# Antigen-dependent inducible T cell reporter system for PET imaging of breast cancer and glioblastoma

**DOI:** 10.1101/2022.02.16.480729

**Authors:** Jaehoon Shin, Matthew F. L. Parker, Iowis Zhu, Aryn Alanizi, Carlos I. Rodriguez, Raymond Liu, Payal B. Watchmaker, Mausam Kalita, Joseph Blecha, Justin Luu, Brian Wright, Suzanne E. Lapi, Robert R. Flavell, Hideho Okada, Thea D. Tlsty, Kole T. Roybal, David M. Wilson

## Abstract

For the past several decades, chimeric antigen receptor T cell (CAR T) therapies have shown promise in the treatment of cancers. These treatments would greatly benefit from companion imaging biomarkers to follow the trafficking of T cells *in vivo*. Using synthetic biology, we engineered T cells with a chimeric receptor SyNthetic Intramembrane Proteolysis Receptor (SNIPR) that induces overexpression of an exogenous reporter gene cassette upon recognition of specific tumor markers. We then applied a SNIPR-based positron emission tomography (PET) reporter system to two cancer-relevant antigens, human epidermal growth factor receptor 2 (HER2) and epidermal growth factor receptor variant III (EGFRvIII), commonly expressed in breast and glial tumors respectively. Antigen-specific reporter induction of the SNIPR-PET T cells was confirmed *in vitro* using GFP fluorescence, luciferase luminescence, and the HSV-TK PET reporter with [^18^F]FHBG. T cells associated with their target antigens were successfully imaged using PET in dual xenograft HER2+/HER2- and EGFRvIII+/EGFRvIII-animal models, with > 10-fold higher [^18^F]FHBG signals seen in antigen-expressing tumors versus the corresponding controls. The main innovation described is therefore PET detection of T cells via specific antigen-induced signals, in contrast to reporter systems relying on constitutive gene expression.

---

Chimeric antigen receptor T cell (CAR T) therapy has revolutionized oncology, demonstrating promising results for refractory drug-resistant leukemias and lymphomas^1–3^ among other cancers. CAR T cells are engineered to respond to cancer cells expressing a specific protein target, inducing rapid cell division and clonal expansion within the tumor microenvironment and activating immune response to target cells via local secretion of cytokines, interleukins, and growth factors^4,5^ **(Supplemental Fig. 1a)**. Hundreds of clinical trials have been initiated globally to study CAR T cells, with two of the most popular targets CD19^6^, and BMCA (seen in multiple myeloma)^7,8^. Importantly, there were engineered T cell clinical trials resulting in patient deaths due to off-target effects, which may have been mediated by recognition of normal lung^9^ or cardiac^10,11^ tissues. We currently do not have tools to detect the engineered T cells engaging antigens in off-target tissues *in vivo* prior to advanced tissue damage, which only can be identified via biopsy or autopsy. Non-invasive methods are therefore critical in evaluating the safety of preclinical CAR T therapies and CAR T clinical trials by providing surrogate real-time maps for patient-specific T cell-target antigen interactions.

Current limitations in predicting CAR T safety *in vivo* are potentially addressed by the positron emission tomography (PET)-compatible SyNthetic Intramembrane Proteolysis Receptor (SNIPR) T cells described in this manuscript. The SNIPR system has produced a powerful new class of chimeric receptors that bind to target surface antigens and induce transcription of exogenous reporter genes via release of a transcription factor domain by regulated intramembrane proteolysis (**Supplemental Fig. 1b**). Importantly, SNIPRs can be designed to have an identical scFv domain as CARs, thereby providing surrogate map for CAR-antigen interaction. The SNIPR contains the regulatory transmembrane domain of human Notch receptor but bears an extracellular antigen recognition domain (eg. single-chain variable fragment or scFv) and an intracellular transcriptional activator domain (Gal4-VP64). When the SNIPR engages its target antigen on an opposing cell, this induces intramembrane cleavage, releasing the intracellular transcriptional domain Gal 4, allowing it to enter the nucleus to activate transcription of target genes^12,13^.

Despite the great versatility of synNotch and SNIPR, their diagnostic potential has not yet been explored. Current PET approaches producing antigen-dependent signals are dominated by immunoPET, whereby a monoclonal antibody is labeled with a radioisotope such as [^89^Zr]^14–16^. In the described SNIPR approach, PET signals also depend on the interaction between an antigen and its corresponding scFv but occur via T cell-based overexpression of the HSV-TK reporter. In this manuscript, we combined CAR and SNIPR technologies to develop a new T cell-based molecular sensor that can image T cells engaged with their target antigens. Upon binding an antigen target, CAR produces rapid T cell division within the tumor microenvironment while SNIPR activates the over-expression of PET-imaging reporter genes. We developed HER2 and EGFRvIII-specific SNIPR T-cells that were successfully imaged *in vivo* upon interaction with their corresponding antigen-expressing tumors. We also compared HER2-specific SNIPR T cells to the “naked” [^89^Zr]-modified anti-HER2 monoclonal antibody ([^89^Zr]herceptin) used in immunoPET to validate the specificity of the cell-based method. This proof-of-concept study provides the foundation for applying the SNIPR-PET reporter to high-sensitivity cell-based antigen detection as well as mapping of engineered T cell-antigen interactions *in vivo*.

## Methods

### Receptor and Response Element Construct Design

#### SNIPR and Chimeric Antigen Receptor (CAR) design

A full description of SNIPR and CAR receptors can be found in the Supplemental Information. SNIPRs were built by fusing anti-HER-2 scFvs or anti-EGFRvIII scFvs with a truncated CD8α hinge region (TTTPAPRPPTPAPTIASQPLSLRPEAC), human Notch1 transmembrane domain (FMYVAAAAFVLLFFVGCGVLLS), intracellular Notch2 juxtamembrane domain (KRKRKH), and a Gal4-VP64 transcriptional element. CARs were built by fusing a binding head (anti-HER2 4D5-8 scFv or IL13 mutein), CD8□ transmembrane domain, co-stimulatory domain 4-1BB, CD3ζ and eGFP.

#### Reporter design

Reporter constructs used are fully described in the Supplemental Information. All reporter constructs were cloned into either a modified pHR’SIN:CSW vector containing a Gal4UAS-RE-CMV promoter followed by multiple cloning site, pGK promoter and mCherry or a modified pHR’SIN:CSW vector containing a Gal4UAS-RE-CMV promoter followed by multiple cloning site, pGK promoter and mCitrine. The HSV-tkSR39-GFP construct was cloned from cEF.tk-GFP (Plasmid #33308, Addgene, MA), which was deposited by Pomper et al.^17^ using site-directed mutagenesis as described in the **Supplemental Fig. 2a**. HSV-tkSR39-T2A-sIL2 construct was cloned from HSV-tkSR39-GFP using In-Fusion cloning (Takara Bio, CA) after adding six C-terminal amino acids (EMGEAN) that were deleted in the original HSV-tk in cEF.tk-GFP, as shown in the **Supplemental Fig. 2b**.

### Preparation of SNIPR T cells

#### Primary Human T cell Isolation and Culture

A full description of T cell isolation and culture is presented in the Supplemental Information. Primary CD4+ and CD8+ T cells were isolated by negative selection (STEMCELL Technologies #15062 & 15063). CD4+ T cells and CD8+ T cells were separated using Biolegend MojoSort Human CD4 T Cell Isolation Kit (Biolegend #480130, San Diego, CA) following manufacturer’s protocol. For experiments involving the induction of Super IL-2, primary T-cells were maintained in human T cell media supplemented with IL-2 until experimentation, whereupon media was replaced with media without added IL-2.

#### Lentiviral Transduction of Human T cells

Transduction of Human T cells was performed as described in the Supplemental Information. Pantropic VSV-G pseudotyped lentivirus was produced via transfection of Lenti-X 293T cells (Clontech #11131D) with a pHR’SIN:CSW transgene expression vector and the viral packaging plasmids pCMVdR8.91 and pMD2.G using Mirus Trans-IT Lenti (Mirus #MIR6606). Primary T cells were thawed the same day, and after 24 hours in culture, were stimulated with Human T-Activator CD3/CD28 Dynabeads (Life Technologies #11131D) at a 1:3 cell:bead ratio. At 48 hours, viral supernatant was harvested, and the primary T cells were exposed to the virus for 24 hours. At day 5 post T cell stimulation, the CD3/CD28 Dynabeads were removed, and the T cells were sorted for assays with a Beckton Dickinson (BD) FACs ARIA II.

#### SNIPR T cell design for in vivo imaging

For HER2+ and HER2-xenograft bioluminescence imaging, we generated anti-HER2 SNIPR T cells with three constructs – constitutively expressed anti-HER2(4D5-8) SNIPR, constitutively expressed anti-HER2(4D5-8) CAR and conditionally expressed firefly luciferase. At baseline, anti-HER2 SNIPR T cells express anti-HER2 SNIPR and anti-HER2 CAR. Upon binding to HER2, they overexpress fLuc through SNIPR activation and proliferate by CAR activation. For HER2+ and HER2-xenograft PET-CT imaging, we generated anti-HER2 SNIPR T cells with three constructs – constitutively expressed anti-HER2(4D5-8) SNIPR, constitutively expressed anti-HER2(4D5-8) CAR and conditionally expressed HSV-tkSR39-T2A-sIL2. At baseline, anti-HER2 SNIPR T cells express anti-HER2 SNIPR and anti-HER2 CAR. Upon binding to HER2, they overexpress HSV-tkSR39 and super IL2 by SNIPR activation, and also proliferate by CAR activation.

For EGFRvIII+ and EGFRvIII-xenograft bioluminescence imaging, we generated anti-EGFRvIII SNIPR T cells with two constructs – constitutively expressed anti-EGFRvIII(139) SNIPR and conditionally expressed IL13 mutein CAR-T2A-nanoLuc. At baseline, anti-EGFRvIII SNIPR T cells express anti-EGFRvIII SNIPR. Upon binding to EGFRvIII, they overexpress IL13 mutein CAR and nanoLuc. For EGFRvIII+ and EVFRvIII-xenograft bioluminescence imaging, we generated anti-EGFRvIII SNIPR T cells with two constructs – constitutively expressed anti-EGFRvIII(139) SNIPR and conditionally expressed anti-IL13m-CAR-HSV-tkSR39. At baseline, anti-EGFRvIII SNIPR T cells express anti-EGFRvIII SNIPR. Upon binding to EGFRvIII, they overexpress IL13 mutein CAR and HSV-tkSR39.

### *In vitro* studies

#### Cancer cell lines

The cell lines used were 293T (ATCC #CRL-3216), MDA-MB-468 (ATCC #HTB-132, HER2-cells), SKBR3 (ATCC # HTB-30, HER2+ high cells), MCF7 (ATCC # HTB-22, HER2+ low cells), U87-EGFRvIII-negative luciferase and U87 EGFRvIII-positive luciferase^12^. Culture media are further described in the Supplemental Information.

#### In vitro fluorophore and luciferase reporter assay

For *in vitro* SNIPR T cell stimulations, 2×10^5^ T cells were co-cultured with 1×10^5^ cancer cells and analyzed at 48-72 hours for reporter expression. Production of fluorophores (GFP and CFP) were assayed using flow cytometry with a BD LSR II and the data were analyzed with FlowJo software (TreeStar). Production of firefly luciferase was assessed with the ONE-glo Luciferase Assay System (Promega #E6110) and production of nanoLuc luciferase was assessed with the Nano-Glo® Luciferase Assay System (Promega #N1110). Bioluminescence was measured with a FlexStation 3 (Molecular Devices).

#### In vitro radiotracer uptake assay

Radiosyntheses of [^18^F]FHBG was performed using established techniques, summarized in **Supplementary Fig. 3** and the supplementary methods for radiosynthesis. For *in vitro* SNIPR T cell stimulations, 1×10^6^ T cells were co-cultured with 5×10^5^ cancer cells and analyzed at 48-72 hours for radiotracer uptake. On the day of radiotracer uptake experiment, T cells and cancer cells were resuspended and 2 μCi of [^18^F]FHBG was added to each well, and incubated for 3 hours at 37°C, 5% CO_2_. After washing, retained radiotracer activity was measured using a Hidex gamma counter (Turku, Finland).

#### Reporter assays with varying receptor affinity and abundance (Heatmap)

Heatmaps for reporter expression were generated as described in the Supplemental Information, with SNIPR receptors of varying HER2 binding affinity (4D5-3, 4D5-5, 4D5-7 and 4D5-8 scFv with the order of increasing binding affinity) and cancer cells with varying amount of surface HER2 expression (293T, MCF7 and SKBR3 with the order of increasing HER2 expression levels).

### *In vivo* studies

#### Murine models/ tumor cohorts studied

Both luciferase-based and PET reporter data were acquired, and two dual xenograft models were studied. After determining the optimal timepoint using optical imaging, (8-10 days), PET imaging was then performed at several time points with sacrifice to verify tissue tracer accumulation (gamma counting) and perform histology and antigen staining^18–20^.

#### Optical Imaging

As shown in **Supplemental Fig. 2**, luciferase-based studies were performed initially (21 days) to investigate the optimal timepoint for [^18^F]FHBG detection of T-cell induction. The SNIPR-T cell distribution within tumor was determined by luminescence emission using a Xenogen IVIS Spectrum after intravenous D-luciferin injection according to the manufacturer’s directions (GoldBio).

#### PET imaging

Radiosyntheses of [^18^F]FHBG, [^18^F]FDG and [^89^Zr]trastzumab: Radiosyntheses of [^18^F]FHBG, [^18^F]FDG and [^89^Zr]trastzumab were performed as described in the Supplemental Methods. Following radiotracer administration, anesthetized under isoflurane and transferred to a Siemens Inveon micro PET-CT system (Siemens, Erlangen, Germany), and imaged using a single static 25 min PET acquisition followed by a 10 min micro-CT scan for attenuation correction and anatomical co-registration.

##### [^18^F]FHBG SNIPR model

Two murine models were studied: (1) a dual HER2+/ HER2-flank model (n = 7) and (2) a dual EGFRvIII+/ EGFRvIII-flank model (n = 4). For the HER2+/HER2– dual xenograft model, First, 1 million MD468 (HER2+ breast cancer cell line, fast growing) and 3 million SKBR3 (HER2-breast cancer cell line, slow growing) subcutaneously injected xenografts generated roughly similar sized tumors in three weeks. Therefore, 4 × 10^6^ SKBR3 cells and 1 × 10^6^ MD468 cells were implanted subcutaneously into 6-10 week-old female NCG mice (Charles River). For the EGFRvIII+/EGFRvIII– dual xenograft model, 1 × 10^6^ EGFRvIII+ or EGFRvIII– U87 cells were implanted subcutaneously into 6-10 week-old female NCG mice (Charles River). All mice were then injected with 6.0 × 10^6^ SNIPR T cells intravenously via tail vein in 100 μl of PBS. For PET imaging, 150 mCi of [^18^F]FHBG was administered via tail vein. At 1 h post-injection, imaging was acquired on day 3, 6, 8, 10 post T cell injection. Upon completion of imaging on day 10, mice were sacrificed, and biodistribution analysis was performed. Gamma counting of harvested tissues was performed using a Hidex Automatic Gamma Counter (Turku, Finland).

##### [^18^F]FDG model

The HER2+/HER2– dual xenograft model was developed as previously described; 4 × 10^6^ SKBR3 cells and 1 × 10^6^ MD468 cells were implanted subcutaneously into 6-10 week-old female NCG mice (Charles River), with 4 mice per group. For PET imaging, 150 uCi of [^18^F]FDG was administered via tail vein. At 1 h post-injection, imaging was acquired.

##### [^89^Zr]trastuzumab model

The same [^18^F]FDG cohort was used (n = 4). For PET imaging, 150 uCi of [^89^Zr]trastuzumab was administered via tail vein. At 3 days post-injection, imaging was acquired. Upon completion of imaging, mice were sacrificed, and biodistribution analysis was performed. Gamma counting of harvested tissues was performed using a Hidex Automatic Gamma Counter (Turku, Finland).

### Data analysis and statistical methods

FACS analysis data were processed using FlowJo (BD biosciences, NJ). For representing data, all graphs are depicted with error bars corresponding to the standard error of the mean. All statistical analysis of *in vitro* data was performed using Microsoft Excel, program language R (https://www.R-project.org/) and Prism software version 7.0 (GraphPad, CA). Data was analyzed using one-way analysis of variance tests (ANOVA) and/or unpaired two-tailed Student’s t-test. PET/μCT data was analyzed using the open source software AMIDE^21^ and %ID/v was used for quantitative comparison. A 95% confidence interval was used to distinguish significant differences in all cases.

## Results

### Synthesis and in vitro validation of PET-compatible antiHER2-SNIPR T cells

To investigate the SNIPR-PET imaging approach, we started with one of the most extensively studied tumor antigens HER2, an extensively studied tumor antigen^22^. Following the published protocol by Roybal et al.^22^, we transduced human CD4+ T cells with two plasmids encoding a SNIPR containing an anti-HER2 scFv (4D5-8), and an inducible reporter. For anti-HER2 SNIPR, we used the SNIPR with an anti-HER2 scFv binding head, an optimized truncated CD8α hinge region, human Notch1 transmembrane domain, intracellular Notch2 juxtamembrane domain, and a transcriptional element composed of GAL4-VP64^23^ (**Fig. 1a**). For the reporter, we used SR39 mutant herpes simplex virus-thymidine kinase (HSV1-sr39tk)-GFP fusion protein and enhanced firefly luciferase (fLuc)^17,24,25^. In this manuscript, we used the hyperactive mutant HSV1-sr39tk to maximize the detection sensitivity^24,26^ (**Supplemental Fig. 2a**). When evaluating tk reporter expression upon SNIPR activation, HSV1-sr39tk-GFP fusion protein was used instead for ready assessment using flow cytometry (**Supplemental Fig. 1b, top**). In all the subsequent radiotracer uptake experiments, however, HSV1-sr39tk was cloned with self-cleavage sequence T2A followed by super IL-2 (sIL-2) instead for higher level of T cell activation upon SNIPR activation (**Supp. Fig. S1b, bottom**).

**Fig. 1.**
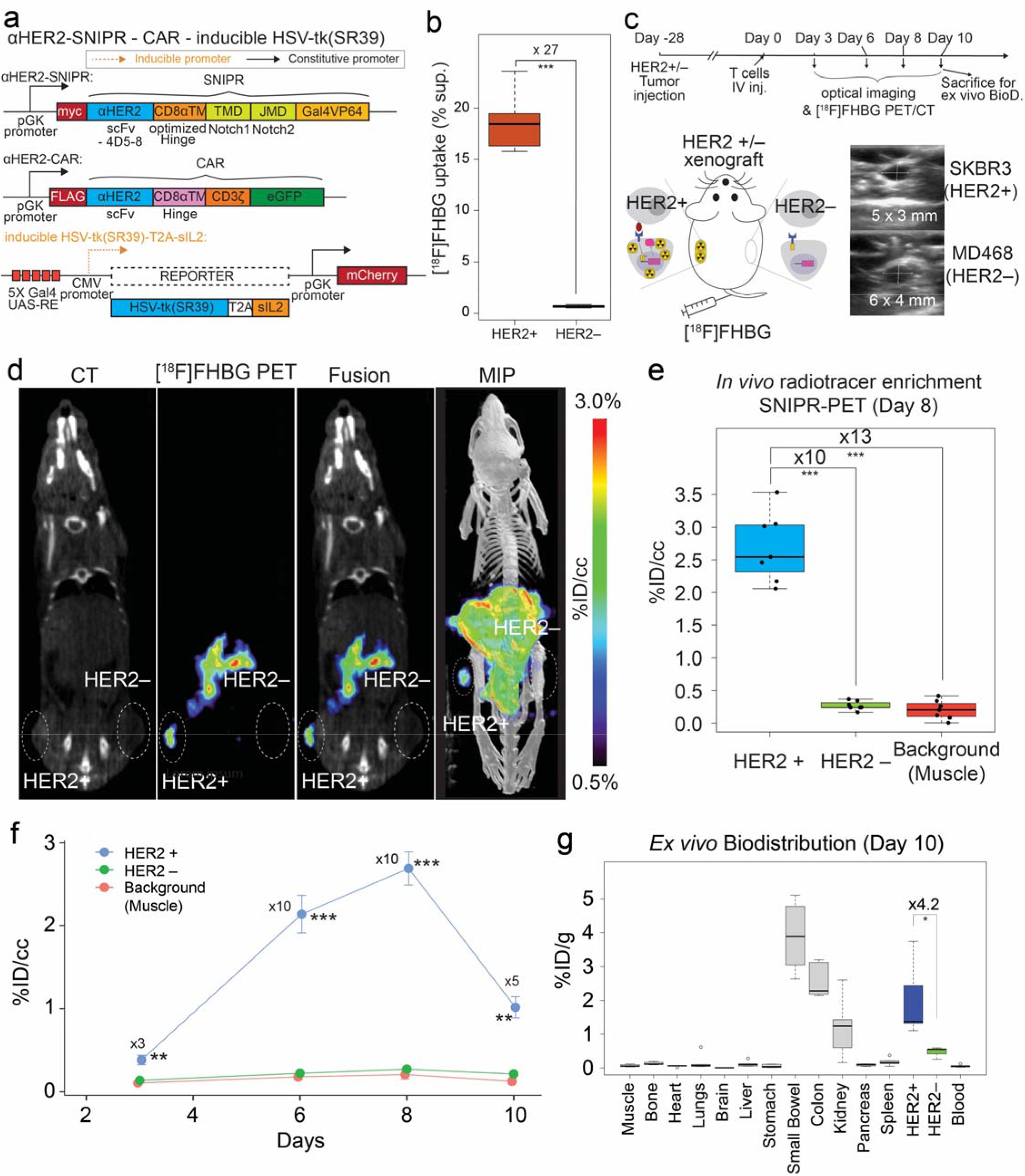
*In vivo* [^18^F]FHBG uptake in HER2+ tumors using activated anti-HER2 SNIPR T cells. a. For PET-CT, we generated T cells by transducing three plasmids including anti-HER2 SNIPR, anti-HER2 CAR and inducible HSV-tk(SR39)-T2A-sIL2, followed by FACS using myc, GFP and mCherry. b. We repeated the *in vitro* radiotracer accumulation of activated SNIPR T cells also bearing CAR, following the experimental scheme shown in the **Fig. 3b**. We confirmed significantly higher [^18^F]FHBG accumulation in SNIPR T cells after co-culturing with SKBR3 (HER2+) cells compared to MD468 (HER2–) cells. c. Double xenograft mouse models were generated by implanting SKBR3(HER2+) and MD468 (HER2–) cells in left and right flank soft tissue. 4 weeks after tumor implantation, SNIPR T cells were injected into tail veins. Micro-PET/CT was acquired 3 days, 6 days, 8 days and 10 days after T cell injection. d. Representative CT images, [^18^F]FHBG PET images, fusion [^18^F]FHBG PET-CT images, and MIP (maximum intensity projection) [^18^F]FHBG PET-CT images at day 8 demonstrate similar size of xenografts with radiotracer enrichment only within the SKBR3 (HER2+) xenograft and not within the MD468 (HER2–) xenograft. e. Quantitative ROI analyses of the HER2+ and HER2– tumors and background (shoulder muscle) at day 8 demonstrate statistically significant radiotracer enrichment within the HER2+ xenograft, 10 times and 13 times greater than within the HER2– xenograft and background. f. Time-dependent ROI analyses of radiotracer enrichment within the HER2+ tumor demonstrate the greatest radiotracer enrichment at day 8 post T cell injection. Slightly decreased radiotracer enrichment was observed at day 10, at which point the mice were sacrificed for *ex vivo* analysis. g. Biodistribution analysis (day 10) of [^18^F]FHBG enrichment within different organs demonstrated significantly greater [^18^F]FHBG enrichment within HER2+ xenograft compared to HER2– xenograft. As seen on microPET-CT, the GI system demonstrated a high level of [^18^F]FHBG uptake. *p<0.05, **p<0.01, ***p<0.001.

We tested the induction of reporters including inducible HSV1-sr39tk-GFP and inducible luciferase activity, in the anti-HER2 SNIPR system (**Supplemental Fig. 4a**). The newly synthesized anti-HER2 SNIPR T cells had comparable transduction efficacy to others we recently published^22^ (**Supplemental Fig. 4b**). Anti-HER2 SNIPR T cells were co-cultured with HER2+ breast cancer line SKBR3 and HER2– breast cancer line MD468 for 48 hours to induce SNIPR activation and downstream reporter gene expression (**Supplemental Fig. 4c**). First, SNIPR activation induced over 160-fold expression of HSV1-sr39tk-GFP compared to negative control (n=4, p < 0.001) (**Supplemental Fig. 4d**). Next, SNIPR activation induced over 30-fold higher luciferase activity compared to negative control (n=4, p<0.001) (**Supplemental Fig. 4e**). We also tested scFvs with varying binding affinities and cancer cells with varying antigen abundance (n = 3 per combination). As expected, HSV1-sr39tk-GFP induction was positively correlated with both SNIPR binding affinity and antigen abundance (**Supplemental Fig. 4f**). The induction of luciferase activity was also positively correlated with the target antigen abundance (n=4, one-way ANOVA p<0.001 for all comparisons of SKBR3)(**Supplemental Fig. 4g**).

### Activated antiHER2-SNIPR T cells show antigen-dependent [^18^F]FHBG accumulation in vitro

Next, we designed SNIPR T cells to secrete super IL-2 (sIL-2), upon binding to target antigens^27^. We created an inducible vector with HSV1-sr39tk and sIL-2, intervened by T2A self-cleaving peptides^28^ (**Supplemental Fig. 2b, bottom, Supplemental Fig. 5a**). We induced the anti-HER2 SNIPR T cells by co-culturing them with SKBR3 or MD468 cells for 48 hours. [^18^F]FHBG was added to the media and SNIPR T cells were incubated for 3 more hours (**Supplmental Fig. 5b**). After removing the supernatant and washing, residual intracellular [^18^F]FHBG was measured using a gamma counter. Anti-HER2 SNIPR T cells co-cultured with SKBR3 cells accumulated over 20-fold higher radiotracer levels compared to anti-HER2 SNIPR T cells co-cultured with MD468 cells (n = 3, p=0.013) (**Supplemental Fig. 5c**). Once again, we tested [^18^F]FHBG radiotracer uptake in suboptimal conditions by using scFvs with lower binding affinities and cancer cells with lower antigen abundance (n = 3 per combination). As expected, [^18^F]FHBG incorporation was correlated with both SNIPR binding affinity and target antigen abundance (**Supplemental Fig. 5d**).

### Imaging using anti-HER2 SNIPR PET shows high antigen specificity in vivo

#### Luciferase-based optical imaging

Since sIL-2 was not strong enough to induce T cell survival and proliferation *in vivo*, we introduced anti-HER2 CAR to the SNIPR T cells to more strongly induce T cell proliferation and survival as well as reporter (HSV1-sr39tk or fLuc) expression upon antigen binding (**Supplemental Fig. 6a**). The FACS yield of T cell transduction with three plasmids was acceptable range, about 10% (**Supplemental Fig. 6b**). In this system, anti-HER2 CAR is constitutively expressed, but only activates T cells and induces proliferation upon binding to its target antigen HER2. As a pilot experiment, we again generated similar sized SKBR3 (HER2+) and MB468 (HER2–) xenografts by subcutaneous injection, followed by anti-HER2 SNIPR T cell injection and bioluminescence imaging for the next 21 days (**Supplemental Fig. 6c**). The luciferase-SNIPR T cell signal was stronger within the HER2+ tumor, which is maximized at day 9, likely reflecting active proliferation within the tumor microenvironment (145-fold, no p value) (**Supplemental Fig. 6d and e**). The signal, however, decreases over time afterwards, likely secondary to minimal target-killing activity of CD4, and clearing of target cells (**Supplemental Fig. 6f**).

#### PET imaging

Based on the luciferase data, we generated SNIPR T cells with constitutively expressed anti-HER2 SNIPR and anti-HER2 CAR and conditionally expressed HSV-sr38tk-T2A-sIL2 (**Fig. 1a**). We first tested the efficacy of *in vitro* [^18^F]FHBG uptake upon SNIPR activation in the SNIPR-CAR system, following the same experimental scheme as the SNIPR only system in **Supplementary Fig. 5B**. As expected, when induced by co-culturing with SKBR3 (HER2+) cells, the SNIPR T cells accumulated 27-times higher amount of [^18^F]FHBG than the SNIPR T cells co-cultured with MD468 (HER2–) cells (N = 8, p<0.001) (**Fig. 1b**). We also chose day 8 to image the SNIPR-T cells with anti-HER2 CAR and inducible HSV1-sr39tk-T2A-sIL2 in HER2+/HER2-xenograft model (**Fig. 1c**). Again, we generated similar sized HER2+(SKBR3) and HER2–(MB468) xenografts by subcutaneous injection into NCG mice, 4 weeks prior to the injection of SNIPR-CAR T cells with HSV-sr38tk-T2A-sIL2 reporter (N=7). [^18^F]FHBG imaging was performed at 3, 6, 8 and 10 days by mPET-CT^29,30^. Greater [^18^F]FHBG uptake on the HER2+ side was observed compared to the HER2-side, indicating that the T cells localized around the HER2+ xenograft (n=7)**(Fig. 1d and e)**. There was an approximately 10-fold higher [^18^F]FHBG uptake on HER2+ xenograft compared to HER2-xenograft (p < 0.001), and approximately 13-fold higher [^18^F]FHBG uptake on HER2+ xenograft compared to shoulder muscle (p < 0.001), based on region of interest (ROI) analysis. In contrast, there was no significant difference in [^18^F]FHBG signal between and background muscle (p > 0.05). As seen on SNIPR-CAR system with luciferase reporter in **Supplemental Fig. 5e**, [^18^F]FHBG signal decreased at 10 days post T cell IV injection, at which time, the tumors were harvested for *ex vivo* biodistribution analysis **(Fig. 1f)**. The difference of PET signal between HER2+ and HER2– xenografts was 5-fold with p=0.003 at day 10, and the difference of ex vivo radioactivity between HER2+ and HER2– xenograft was 4.2-fold with p=0.036 (**Fig. 1f, g**). Images from the PET study showed marked radiotracer in the stomach, intestine, and gallbladder consistent with the known biodistribution of [^18^F]FHBG in wild type animals due to hepatobiliary excretion, as shown in both human and rodent subjects (**Fig. 3d**)^31–33^. This result was corroborated using tissue extraction and *ex vivo* gamma-counting at day 10 (**Fig. 1g**).

### Comparing anti-HER2 SNIPR-PET with anti-HER2 [^89^Zr]trastuzumab and [^18^F]FDG

Trastuzumab has the same anti-HER2 scFv binding moiety as our SNIPR, thereby reflecting the affinity-based interaction of the same antigen/antibody pair^34^. We used [^89^Zr]trastuzumab (anti-HER2) PET-imaging as well as [^18^F]FDG-PET in the same animal model as SNIPR-CAR system (N = 4). Overall, different biodistributions of the two tracers were observed, consistent with distinct metabolism and excretion pathways (**Fig. 2a**). Both immunoPET with [^89^Zr]trastuzumab and SNIPR-PET with [^18^F]FHBG demonstrated statistically significant increased radiotracer enrichment in HER2+ tumor compared to HER2– tumor (9.9-fold with p<0.001 and 9.3-fold with p=0.002, respectively)(**Fig. 2b**). The relative radiotracer enrichment within HER2+ tumor compared to HER2-tumor was not statistically significant between ImmunoPET and SNIPR-PET (p > 0.05) (**Fig. 2c**). Likewise, the relative radiotracer enrichment within HER2+ tumor compared to background was also not statistically significant between ImmunoPET and SNIPR-PET (p > 0.05) (**Fig. 2d**). Imaging results using [^89^Zr]trastuzumab were corroborated via *ex vivo* analysis of harvested tissues (**Supplemental Fig. 7**). Although not statistically significant, the trend of higher [^18^F]FDG accumulation in HER2– tumor compared to HER2+ tumor on a %ID/cc basis correlated with the higher growth rate of MD468 (HER2–) compared to SKBR3 (HER2+) we observed both *in vitro* and *in vivo* (**Fig. 2b, middle and 2d**).

**Fig. 2.**
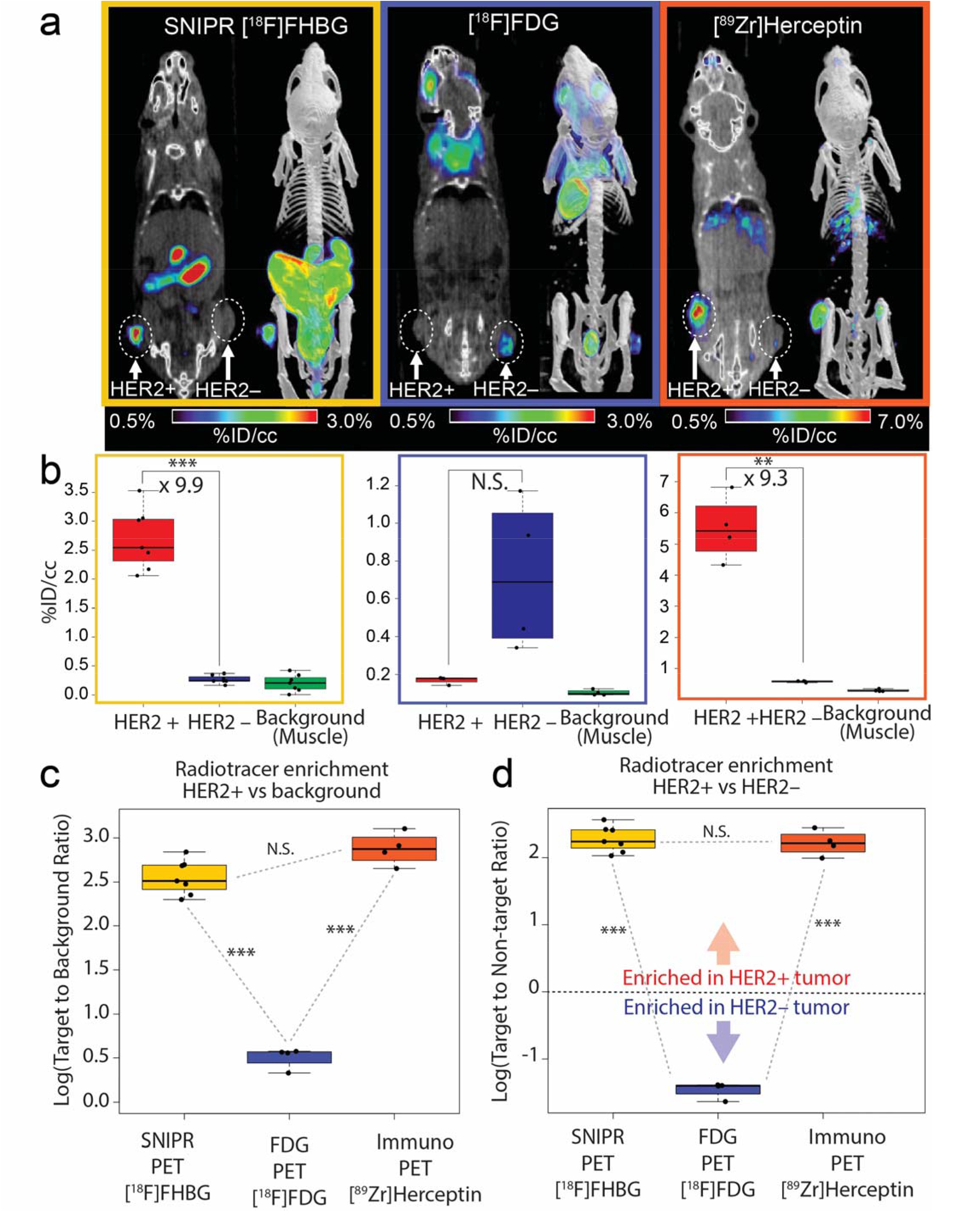
Comparison between SNIPR-PET, FDG-PET and ImmunoPET. a. Representative images of [^18^F]FHBG SNIPR-PET, [^18^F]FDG-PET and [^89^Zr]-herceptin-PET. SNIPR-PET and Herceptin immunoPET demonstrated radiotracer enrichment within SKBR3(HER2+) xenograft compared to MD468 (HER2–) tumor, whereas FDG-PET demonstrated higher radiotracer enrichment within MD468 tumor. b. ROI analysis demonstrated statistically significant, 9.9-fold greater enrichment of [^18^F]FHBG within HER2+ tumor than within HER2– tumor (left), non-statistically significant enrichment (p-value > 0.05) of FDG within HER2– tumor compared to HER2+ tumor, and statistically significant 9.3 times greater enrichment of [^89^Zr]Herceptin within HER2+ tumor than within HER2– tumor. c and d. Fold enrichment of radiotracer within HER2+ tumor was significantly greater in SNIPR-PET and ImmunoPET compared to FDG-PET, when compared to background (c) and when compared to HER2– tumor (d). *p<0.05, **p<0.01, ***p<0.001.

### SNIPR_PET can be extended to the glioblastoma antigen EGFRvIII

To demonstrate the feasibility of SNIPR-PET in other cancers, we chose the glioblastoma-specific antigen EGFRvIII. We designed T cells to constitutively express SNIPR that targets EGFRvIII and conditionally express CAR that targets distinct antigen, IL13Rα2, analogous to our previous approach for targeting glioblastoma with cytotoxic CD8 T cells^12^. At baseline, those T cells express only anti-EGFRvIII SNIPR, without CAR or reporter. When they recognize EGFRvIII, they express IL13-mutein (IL13m)-CAR that strongly binds to more widely expressed but less specific target interleukin 13 receptor alpha 2 (IL13Rα2), as well as reporter genes (**Fig. 3a**). To demonstrate *in vitro* reporter activation upon anti-EGFRvIII SNIPR activation, we generated three different T cells with Blue Fluorescent Protein (BFP), nano-Luciferase (nLuc) and tkSR39 reporters. Those reporters were co-expressed with IL13m-CAR, and then cleaved at the intervening T2A sequence to generate separate CAR and reporter proteins. As expected, T cells bearing anti-EGFRvIII SNIPR receptor induced a significantly higher level of BFP and nLuc activity 48 hours after co-culturing with EGFRvIII+ U87 cells, compared to co-culturing with EGFRvIII– U87 cells (BFP: n=4, 25-fold, p<0.001; nLuc: n=4, p=0.002) (**Fig. 3b, left and middle**). T cells bearing anti-EGFRvIII SNIPR receptor and inducible IL13m-CAR-T2A-tkSR39 demonstrated a significantly higher level of [^18^F]FHBG uptake when co-cultured with EGFRvIII+ U87 cells compared to the same T cells co-cultured with EGFRvIII– U87 cells (n=8, 32.5-fold, p<0.001). To demonstrate the potential for *in vivo* imaging, we again generated a mouse model with EGFRvIII+ U87 and EGFRvIII– U87 xenografts by subcutaneous injection, followed by SNIPR T cell injection and PET imaging for the next 10 days (n = 4) (**Fig. 3c**). ROI analysis demonstrated a significantly higher level of radiotracer within EGFRvIII+ xenograft compared to EGFRvIII– xenograft, which was maximized at day 8, based on per volume radioactivity (%ID/cc) (14-fold, p<0.001) (**Fig. 3d and e**). As seen on the HER2 SNIPR-CAR system, EGFRvIII SNIPR CAR system demonstrated a decrease in PET signal at day 10, likely secondary to low target killing activity of activated CD4+ T cells (**Fig. 3d)**. Again, EGFRvIII+ and EGFRvIII-xenografts were sacrificed at day 10 for ex vivo analysis, which confirmed significantly higher enrichment of radiotracer within the EGFRvIII+ xenograft compared to EGFRvIII– xenograft based on per weight radioactivity (%ID/g) 5-fold excess, p<0.001) (**Fig. 3f)**.

**Fig. 3.**
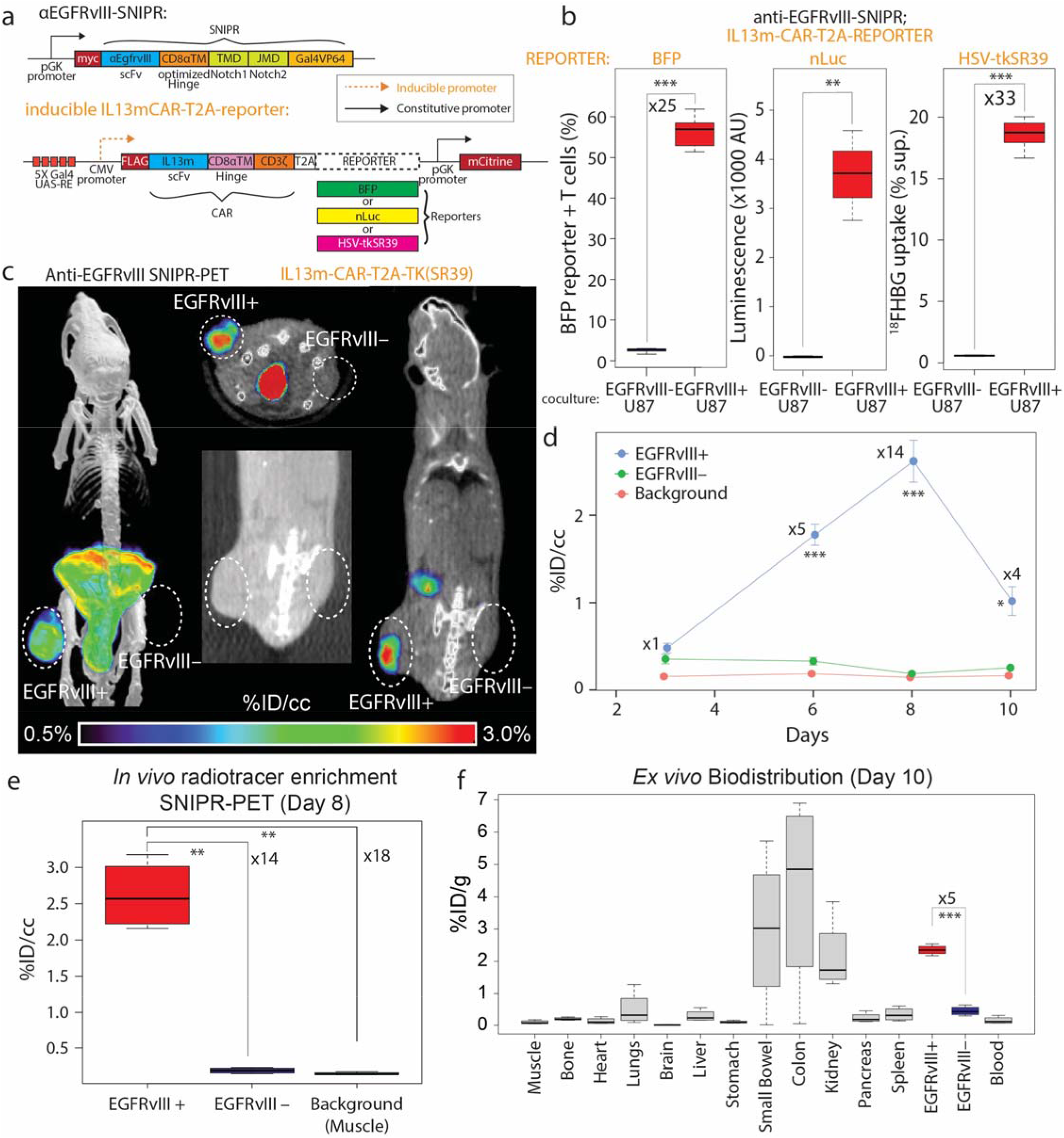
EGFRvIII SNIPR-PET. a. We generated SNIPR-T cells with anti-EGFRvIII-SNIPR and inducible IL13-mutein (IL13m)-CAR-T2A-reporter constructs. We used three different reporters – Blue Fluorescent Protein (BFP), nanoLuc (nLuc) and HSV-tkSR39. In this system, SNIPR T cells express anti-EGFRvIII-SNIPR at baseline but do not express IL13m-CAR or reporters. When anti-EGFRvIII binds to EGFRvIII on target cells, SNIPR T cells now induce the expression of IL13m-CAR and reporters - BFP, nLuc or HSV-tkSR39. Since most U87 cells express IL13Rα2, T cells expressing IL13m-CAR now secrete cytokines and growth factors that induce T cell proliferation and survival. b. SNIPR T cells incubated with EGFRvIII+ U87 cells demonstrated significantly higher level of BFP reporter expression, significantly greater level of nLuc enzymatic activity and HSV-tkSR39-mediated ^18^FHBG accumulation compared to SNIPR T cells incubated with EGFRvIII– U87 cells. c. Following a similar protocol as used for HER2, EGFRvIII+ U87 and EGFRvIII– U87 cells were implanted into mouse flank subcutaneous tissues. 4 weeks after implantation, anti-EGFRvIII T cells with inducible anti-IL13-mutein-CAR-T2A-HSV-tk(SR39) were injected into the tail veins. Representative MIP [^18^F]FHBG PET-CT image (left) and cross-sectional [^18^F]FHBG PET-CT images (middle and right) at day 8 demonstrate high radiotracer enrichment within the EGFRvIII+ U87 xenograft compared to the EGFRvIII– U87 xenograft on the contra-lateral side. d. Time dependent ROI analysis of radiotracer enrichment within the EGFRvIII+ xenograft demonstrates the greatest radiotracer enrichment at day 8 post T cell injection, followed by slight decrease in PET signal at day 10, at which point, animals were sacrificed for ex vivo biodistribution analysis. e. Quantitative ROI analysis of the EGFRvIII+ and EGFRvIII– tumors and background (shoulder muscle) demonstrate statistically significant radiotracer enrichment within the EGFRvIII+ xenograft, 14 times and 18 times greater than within the EGFRvIII– xenograft and background. f. *Ex vivo* analysis (day 10) of [^18^F]FHBG enrichment within different organs demonstrated significantly greater [^18^F]FHBG enrichment within EGFRvIII+ xenograft compared to EGFRvIII– xenograft. As seen on microPET-CT images, the GI system demonstrated a high level of [^18^F]FHBG. *p<0.05, **p<0.01, ***p<0.001.

## Discussion

We have developed an antigen-specific, PET-compatible SNIPR T cell reporter system, in response to the rapidly proliferating interest in T cell-mediated treatment of human tumors. This customizable synthetic receptor platform can provide an essential companion to CAR T mediated treatment, in addition to the recent application of synNotch to oncologic challenges^12^. Applying several rounds of synthetic biology, we engineered SNIPR T cells that (1) are antigen-inducible, as detected by GFP fluorescence, luciferase luminescence, and the HSV-TK PET reporter [^18^F]FHBG and (2) can be successfully imaged dual xenograft animal models. We have demonstrated application of this imaging modality to two different tumor antigens-HER2 and EGFRvIII. This technology could be used as a companion biomarker for CAR T therapies, for example to characterize off-target effects, or verify tumor engagement. As the technology matures, SNIPR-PET could be used to detect and characterize molecular profiles of early cancers without biopsy.

## Key Points

*Question:* Can an inducible, cell-based PET system image tumor antigens *in vivo*?

*Pertinent findings:* An inducible PET reporter system can be co-expressed with CAR in therapeutic T cells to image HER2 and EGFRvIII.

*Implications for patient care:* As cell-based therapies mature, SNIPR-PET could be added to any therapeutic T cell to image antigen engagement.

## Acknowledgements

Grant sponsors NIH (R01-EB024014, R01-EB025985); Kleberg Foundation 132472B; Society of Interventional Radiology.

## Author contributions

J.S., K.T.R., D.M.W. and T.D.T. proposed and supervised the overall project. J.B. performed or supported the radiochemistry. J.S., I.Z., R.L., and C.I.R. generated the receptor and reporter constructs and generated SNIPR T cells. P.B.W. and H.O. generated EGFRvIII+ and EGFRvIII-cell lines. M.F.L.P. generated HER2 and EGFRvIII xenografts. M.F.L.P., A.A., J.L. and M.K. performed the *in vitro* PET studies. M.F.L.P. and A.A. performed µPET-CT imaging studies and J.S., M.F.L.P. and R.F. performed subsequent data analysis. M.F.L.P. and A.A. performed *ex vivo* analysis. C.I.R. and T.D.T performed histology analysis of *ex vivo* specimens. J.S., K.T.R. and D.M.W. wrote and edited the paper.

## Materials and Correspondence

*
*
*

## Supplementary information

Please see the supplementary information for detailed information regarding molecular biology, radiosynthesis, and several imaging studies not reported in the main text.

## For Table of Contents use only

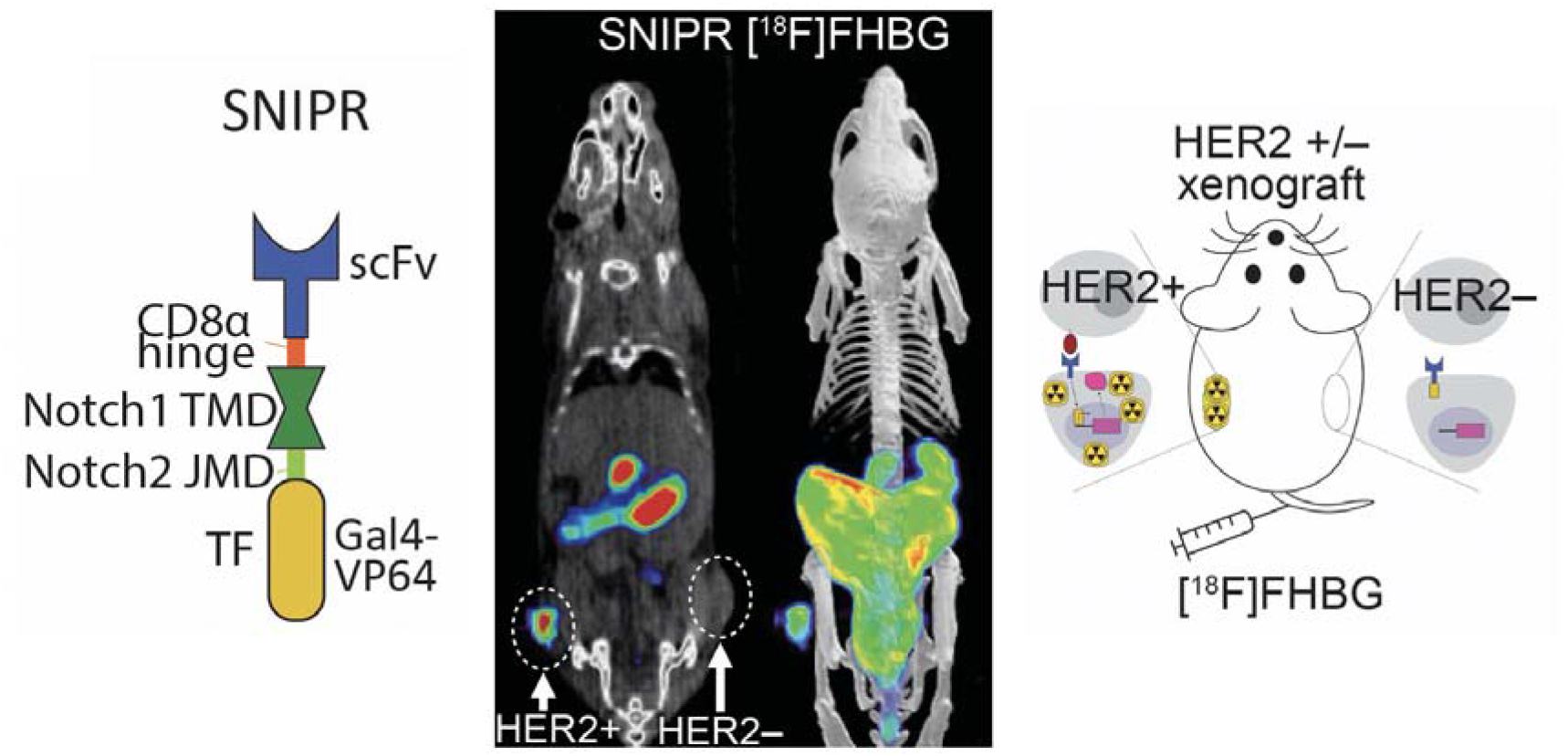

