## Supplementary material for "Antigen-dependent inducible T cell reporter system for PET imaging of breast cancer and glioblastoma": SI

[1] Department of Radiology and Biomedical Imaging, University of California, San Francisco, San Francisco, CA, USA
[2] Department of Microbiology and Immunology, University of California, San Francisco, San Francisco, CA, USA
[3] Parker Institute for Cancer Immunotherapy, San Francisco, CA, USA
[4] Department of Pathology, University of California, San Francisco, San Francisco, CA, USA
[5] Department of Neurological Surgery, University of California, San Francisco, CA, USA
[6] Department of Radiology, University of Alabama at Birmingham, AL, USA
[7] Helen Diller Cancer Center, University of California, San Francisco, San Francisco, CA, USA.
[8] Chan Zuckerberg Biohub, San Francisco, CA, USA
[9] Gladstone UCSF Institute for Genetic Immunology, San Francisco, CA, USA
[10] UCSF Cell Design Institute, San Francisco, CA, USA
[11] Current address: Department of Psychiatry, Renaissance School of Medicine at Stony Brook University, Stony Brook, NY, USA
[12] These authors contributed equally: Jaehoon Shin, Matthew F. L. Parker, Iowis Zhu
✉

---

**SUPPLEMENTARY METHODS:**

**Receptor and Response Element Construct Design**

*SNIPR receptor design*

SNIPRs were built by fusing anti-HER-2 scFvs or anti-EGFRvIII scFvs with a truncated CD8α hinge region (TTTPAPRPPTPAPTIASQPLSLRPEAC), human Notch1 transmembrane domain (FMYVAAAAFVLLFFVGCGVLLS), intracellular Notch2 juxtamembrane domain (KRKRKH), and a Gal4-VP64 transcriptional element. All receptors contain an N-terminal CD8α signal peptide (MALPVTALLLPLALLLHAARP) for membrane targeting and a myc-tag (EQKLISEEDL) for easy determination of surface expression with α-myc AF647 (Cell-Signaling #2233). Four versions of antiHER-2 scFv with different affinities, 4D5-3, 4D5-5, 4D5-7 and 4D5-8[1], and two versions of anti-EGFRviii scFv, 139 and 3C10[2], were tested and 4D5-8 for HER2 system and 139 for EGFRvIII system were used if not otherwise stated. Receptors were cloned into a modified pHR'SIN:CSW vector containing a PGK promoter for all primary T cell experiments.

*Chimeric Antigen Receptor (CAR) design*

CARs were built by fusing a binding head (anti-HER2 4D5-8 scFv or IL13 mutein), CD8α transmembrane domain, , co-stimulatory domain 4-1BB, CD3ζ and eGFP. Of note, eGFP in the C terminal of CAR did not affect the function of CAR. All receptors contain an N-terminal CD8α signal peptide (MALPVTALLLPLALLLHAARP) for membrane targeting and FLAG tag (DYKDDDDK) for easy determination of surface expression. The anti-HER2 CAR was cloned into a modified pHR'SIN:CSW vector containing a constitutive pGK promoter for *in vivo* experiments. The IL13-mutein CAR was cloned as inducible vector by cloning into pHR'SIN:CSW that contains five copies of the Gal4 DNA binding domain target sequence (GGAGCACTGTCCTCCGAACG), followed by an inducible CMV promoter. In inducible vectors, IL13 mutein CAR was followed by T2A-HSV-tkSR39 or T2A-nLuc for co-induction with reporter genes. All inducible CAR-T2A-reporter vectors contained constitutively expressed fluorophore genes (mCitrine or mCherry) to easily identify transduced T cells. All induced elements were cloned via a BamHI site in the multiple cloning site 3′ to the Gal4 response elements. All constructs were cloned via In-Fusion cloning (Takara # 638951).

*Reporter design*

All reporter constructs were cloned into either a modified pHR'SIN:CSW vector containing a Gal4UAS-RE-CMV promoter followed by multiple cloning site, pGK promoter and mCherry or a modified pHR'SIN:CSW vector containing a Gal4UAS-RE-CMV promoter followed by multiple cloning site, pGK promoter and mCitrine. All reporter-T2A-sIL2 constructs were cloned using pre-existing sIL2 constructs in the lab, stem from pre-existing constructs containing sIL2, based on Levine et al.[3]. HSV-tkSR39-GFP construct was cloned from cEF.tk-GFP (Plasmid #33308, Addgene, MA), which was deposited by Pomper et al.[4] using site-directed mutagenesis as described in the **Supplemental Fig. 2A**. HSV-tkSR39-T2A-sIL2 construct was cloned from HSV-tkSR39-GFP using In-Fusion cloning (Takara Bio, CA) after adding six C-terminal amino acids (EMGEAN) that were deleted in the original HSV-tk in cEF.tk-GFP, as shown in the **Supplemental Fig. 2 A,B**, nLuc-T2A-sIL2 was cloned from pPRE-pNL1.3 (Plasmid #84394, Addgene, MA). Inducible IL13m-CAR-T2A-reporter constructs were cloned as described above.

### Preparation of SNIPR T cells:

*Primary Human T cell Isolation and Culture*

Primary CD4+ and CD8+ T cells were isolated from anonymous donor blood after apheresis by negative selection (STEMCELL Technologies #15062 & 15063). Blood was obtained from Blood Centers of the Pacific (San Francisco, CA) as approved by the University Institutional Review Board. CD4+ T cells and CD8+ T cells were separated using Biolegend MojoSort Human CD4 T Cell Isolation Kit (Biolegend #480130, San Diego, CA) following manufacturer's protocol. In all experiments in this manuscript, we used human CD4+ T cells with minimal target-killing activity. T cells were cryopreserved in RPMI-1640 (Thermo Fisher #11875093) with 20% human AB serum (Valley Biomedical Inc., #HP1022) and 10% DMSO. After thawing, T cells were cultured in human T cell medium consisting of X- VIVO 15 (Lonza #04-418Q), 5% Human AB serum and 10 mM neutralized N-acetyl L- Cysteine (Sigma-Aldrich #A9165) supplemented with 30 units/mL IL-2 (NCI BRB Preclinical Repository) for most experiments. For experiments involving the induction of Super IL-2, primary T-cells were maintained in human T cell media supplemented with IL-2 until experimentation, whereupon media was replaced with media without supplemented IL-2.

*Lentiviral Transduction of Human T cells*

Pantropic VSV-G pseudotyped lentivirus was produced via transfection of Lenti-X 293T cells (Clontech #11131D) with a pHR'SIN:CSW transgene expression vector and the viral packaging plasmids pCMVdR8.91 and pMD2.G using Mirus Trans-IT Lenti (Mirus #MIR6606). Primary T cells were thawed the same day, and after 24 hours in culture, were stimulated with Human T-Activator CD3/CD28 Dynabeads (Life Technologies #11131D) at a 1:3 cell:bead ratio. At 48 hours, viral supernatant was harvested, and the primary T cells were exposed to the virus for 24 hours. At day 5 post T cell stimulation, the CD3/CD28 Dynabeads were removed, and the T cells were sorted for assays with a Beckton Dickinson (BD) FACs ARIA II. Sorted T-cells were cultured in the T cell media for 24 hours, CD3/CD28 Dynabeads was added at a 1:3 cell:bead ratio to expand for the next 3 days. CD3/CD28 Dynabeads were removed, and the cells were further cultured/expanded for 7 days, at which point, almost all T cells stopped dividing. Those T cells were cryopreserved in RPMI-1640 (Thermo Fisher #11875093) with 20% human AB serum (Valley Biomedical Inc., #HP1022) and 10% DMSO. 7 days before *in vitro* or *in vivo* experiments, the cryopreserved T cells were thawed and cultured in T cell media with CD3/CD28 Dynabeads at a 1:3 cell:bead ratio for the first 3 days, and the beads were removed, and T cells were cultured 4 additional days to use in experiments.

*SNIPR T cell design for in vivo imaging*

For HER2+ and HER2- xenograft bioluminescence imaging, we generated anti-HER2 SNIPR T cells with three constructs – constitutively expressed anti-HER2(4D5-8) SNIPR, constitutively expressed anti-HER2(4D5-8) CAR and conditionally expressed firefly luciferase (**Supplemental Fig. 4A**). At baseline, anti-HER2 SNIPR T cells express anti-HER2 SNIPR and anti-HER2 CAR. Upon binding to HER2, they overexpress fLuc through SNIPR activation and proliferate by CAR activation. For HER2+ and HER2- xenograft PET-CT imaging, we generated anti-HER2 SNIPR T cells with three constructs – constitutively expressed anti-HER2(4D5-8) SNIPR, constitutively expressed anti-HER2(4D5-8) CAR and conditionally expressed HSV-tkSR39-T2A-sIL2 (**Fig. 1A**).

At baseline, anti-HER2 SNIPR T cells express anti-HER2 SNIPR and anti-HER2 CAR. Upon binding to HER2, they overexpress HSV-tkSR39 and super IL2 by SNIPR activation, and also proliferate by CAR activation.

For EGFRvIII+ and EGFRvIII- xenograft bioluminescence imaging, we generated anti-EGFRvIII SNIPR T cells with two constructs – constitutively expressed anti-EGFRvIII(139) SNIPR and conditionally expressed IL13 mutein CAR-T2A-nanoLuc (**Fig. 3A**). At baseline, anti-EGFRvIII SNIPR T cells express anti-EGFRvIII SNIPR. Upon binding to EGFRvIII, they overexpress IL13 mutein CAR and nanoLuc. For EGFRvIII+ and EVFRvIII- xenograft bioluminescence imaging, we generated anti-EGFRvIII SNIPR T cells with two constructs – constitutively expressed anti-EGFRvIII(139) SNIPR and conditionally expressed anti-IL13m-CAR-HSV-tkSR39. At baseline, anti-EGFRvIII SNIPR T cells express anti-EGFRvIII SNIPR. Upon binding to EGFRvIII, they overexpress IL13 mutein CAR and HSV-tkSR39.

***In vitro* studies:**

*Cancer cell lines*

The cell lines used were 293T (ATCC #CRL-3216), MDA-MB-468 (ATCC #HTB-132, HER2- cells), SKBR3 (ATCC # HTB-30, HER2+ high cells), MCF7 (ATCC # HTB-22, HER2+ low cells), U87-EGFRvIII-negative luciferase and U87 EGFRvIII-positive luciferase (Choe et al., 2021). All cell lines in this manuscript were cultured in filter sterilized DMEM (Sigma D5648) with 10% FBS and 1% Penicillin/Streptomycin stock (ThermoFisher, 10,000 U penicillin and 10,000 µg/ml streptomycin).

*In vitro fluorophore and luciferase reporter assay*

For *in vitro* SNIPR T cell stimulations, $2x10^5$ T cells were co-cultured with $1x10^5$ cancer cells. After mixing the T cells and cancer cells in flat bottom 96-well tissue culture plates, the cells were centrifuged for 1 min at 400xg to force interaction of the cells and the cultures were analyzed at 48-72 hours for reporter expression. Production of fluorophores (GFP and CFP) were assayed using flow cytometry with a BD LSR II and the data were analyzed with FlowJo software (TreeStar). Production of firefly luciferase was assessed with the ONE-glo Luciferase Assay System (Promega #E6110) and production of nanoLuc luciferase was assessed with the Nano-Glo®

Luciferase Assay System (Promega #N1110). Bioluminescence was measured with a FlexStation 3 (Molecular Devices).

*In vitro radiotracer uptake assay*

For *in vitro* SNIPR T cell stimulations, $1 \times 10^6$ T cells were co-cultured with $5 \times 10^5$ cancer cells. After mixing the T cells and cancer cells in 6-well tissue culture plates, the cells were centrifuged for 1 min at 400xg to force interaction of the cells and the cultures were analyzed at 48-72 hours for radiotracer uptake. On the day of radiotracer uptake experiment, T cells and cancer cells were resuspended and 2 µCi of [$^{18}$F]FHBG was added to each well, and incubated for 3 hours at 37˚C, 5% $CO_2$. Cells were resuspended and washed three times with 4˚C 2% FBS PBS. The retained radiotracer activity was measured using a Hidex gamma counter (Turku, Finland).

*Reporter assays with varying receptor affinity and abundance (Heatmap)*

The heatmap for reporter expression was generated with four versions of anti-HER2 SNIPR T cells bearing anti-HER2 SNIPR receptor with varying HER2 binding affinity (4D5-3, 4D5-5, 4D5-7 and 4D5-8 scFv with the order of increasing binding affinity). All the SNIPR T cells had inducible reporter of HSV-tk-GFP fusion protein that is to be detected with flow cytometry. SNIPR T cells were incubated with cancer cells with varying amount of surface HER2 expression (293T, MD468, MCF7 and SKBR3 with the order of increasing HER2 expression levels). $2 \times 10^5$ SNIPR T cells were co-cultured with $1 \times 10^5$ cancer cells in flat bottom 96-well tissue culture plates for 48-72 hours and the production of HSV-tk-GFP was assayed using flow cytometry with a BD LSR II and the data were analyzed with FlowJo software (TreeStar).

The heatmap for radiotracer accumulation was generated with three versions of anti-HER2 SNIPR T cells bearing anti-HER2 SNIPR receptor with varying HER2 binding affinity (4D5-5, 4D5-7 and 4D5-8 scFv). All the SNIPR T cells had reporter of HSV-tk-T2A-sIL2 protein. SNIPR T cells were incubated with cancer cells with varying amount of surface HER2 expression (293T, MCF7 and SKBR3 with the order of increasing HER2 expression levels). $1 \times 10^6$ SNIPR T cells were co-cultured with $5 \times 10^5$ cancer cells in flat bottom 6-well tissue culture plates for 48-72 hours. The FHBG radiotracer uptake was measured using a Hidex gamma counter.

***In vivo* studies:**

*Murine models/ tumor cohorts studied:*

All mouse experiments were conducted according to Institutional Animal Care and Use Committee (IACUC)–approved protocols. Both luciferase-based and PET reporter data were acquired, and two dual xenograft models were studied. After determining the optimal timepoint using optical imaging, (8-10 days), PET imaging was then performed at several time points with sacrifice to verify tissue tracer accumulation (gamma counting) and perform histology and antigen staining[5–7].

*Optical Imaging:*

As shown in **Supplemental Fig. 6**, luciferase-based studies were performed initially (21 days) to investigate the optimal timepoint for [$^{18}$F]FHBG detection of T-cell induction. The SNIPR-T cell distribution within tumor was determined by luminescence emission using a Xenogen IVIS Spectrum after intravenous D-luciferin injection according to the manufacturer's directions (GoldBio).

*PET imaging:*

Radiosyntheses of [$^{18}$F]FHBG, [$^{18}$F]FDG and [$^{89}$Zr]trastzumab: Radiosyntheses of [$^{18}$F]FHBG, [$^{18}$F]FDG and [$^{89}$Zr]trastzumab were performed as described in the supplementary methods.

Imaging protocol: The same general imaging protocol was used for all studies. A tail vein catheter was placed in mice under isoflurane anesthesia and the radio tracer was subsequently injected. The animals were placed on a heating pad to minimize shivering. Mice were allowed to recover and micturate, and at specific later time point, the animals were again anesthetized under isoflurane and transferred to a Siemens Inveon micro PET-CT system (Siemens, Erlangen, Germany), and imaged using a single static 25 min PET acquisition followed by a 10 min micro-CT scan for attenuation correction and anatomical co-registration. No adverse events were observed during or after injection of any compound. Anesthesia was maintained during imaging using isofluorane.

*[$^{18}$F]FHBG SNIPR model:* Two murine models were studied: (1) a dual HER2+/ HER2- flank model (n = 7) and (2) a dual EGFRvIII+/ EGFRvIII- flank model (n = 4). For the HER2+/HER2– dual xenograft model, First, 1 million MD468 (HER2+ breast cancer cell line, fast growing) and 3 million SKBR3 (HER2- breast cancer cell line, slow growing) subcutaneously injected xenografts generated roughly similar sized tumors in three weeks. Therefore, 4 x $10^6$ SKBR3 cells and 1 x $10^6$ MD468 cells were implanted subcutaneously into 6-10 week-old female NCG mice (Charles River). For the EGFRvIII+/EGFRvIII– dual xenograft model, 1 x $10^6$ EGFRvIII+ or EGFRvIII– U87 cells were implanted subcutaneously into 6-10 week-old female NCG mice (Charles River). All mice were then injected with 6.0 × $10^6$ SNIPR T cells intravenously via tail vein in 100 µl of PBS. For PET imaging, 150 mCi of [$^{18}$F]FHBG was administered via tail vein. At 1 h post-injection, imaging was acquired on day 3, 6, 8, 10 post T cell injection. Upon completion of imaging on day 10, mice were sacrificed, and biodistribution analysis was performed. Gamma counting of harvested tissues was performed using a Hidex Automatic Gamma Counter (Turku, Finland).

*[$^{18}$F]FDG model:* The HER2+/HER2– dual xenograft model was developed as previously described; 4 x $10^6$ SKBR3 cells and 1 x $10^6$ MD468 cells were implanted subcutaneously into 6-10 week-old female NCG mice (Charles River), with 4 mice per group. For PET imaging, 150 uCi of [$^{18}$F]FDG was administered via tail vein. At 1 h post-injection, imaging was acquired.

*[$^{89}$Zr]trastuzumab model:* The same [$^{18}$F]FDG cohort was used (n = 4). For PET imaging, 150 uCi of [$^{89}$Zr]trastuzumab was administered via tail vein. At 3 days post-injection, imaging was acquired. Upon completion of imaging, mice were sacrificed, and biodistribution analysis was performed. Gamma counting of harvested tissues was performed using a Hidex Automatic Gamma Counter (Turku, Finland).

**Radiotracer syntheses:**

*Radiosynthesis of [$^{18}$F]FHBG:*

[$^{18}$F]FHBG was synthesized according to a previously published method[8], as shown in **Supplemental Fig. 3**.

In short, [$^{18}$F]FHBG was synthesized on the ELIXYS FLEX/ CHEM (Sofie Biosciences, VA) from [$^{18}$F]fluoride ion. The [$^{18}$F]fluoride ion was trapped on a QMA cartridge (Waters, MA) and eluted with $K_2CO_3$/ K222 (4.75 mg/ 25 mg) in acetonitrile/ water. The resultant solution was azeotroped at 110 ˚C twice with additions of acetonitrile. To this dried vial the precursor (ABX, Radeberg, Germany) dissolved in acetonitrile (0.8 mL) was added and heated to 140 ˚C for 10 minutes. Then 1 mL of 1M HCl was added and the reactor was heated to 105 ˚C for 5 minutes. The reaction was cooled, neutralized with 0.5 mL of 2 M NaOH, diluted with solvent system (2 mL), and injected onto the semi-preparative HPLC (C18(2) 250 x 10 mm, 7% EtOH/ 93% 50 mM $NH_4OAc$, 5 mL/ min, 254 nm). The desired product was collected, filtered and analyzed via HPLC (C18(2) 250 x 4.6 mm, 7% EtOH/ 93% 50 mM NH4OAc, 1 mL/ min, 254 nm). [$^{18}$F]FHBG was synthesized with a decay corrected yield of 11.1 +/- 3.9 % in 68.3 +/- 5.4 minutes ( n = 18). The radiochemical purity of the synthesized [$^{18}$F]FHBG was > 99%.

*Radiosynthesis of $^{89}$Z-Trastuzumab:*

Trastuzumab-DFO was purchased from Dr. Suzanne E. Lapi from University of Alabama at Birmingham. Please see Chang et al. for a detailed protocol[9]. Briefly, Trastuzumab (21 mg/mL) was conjugated to DFO-Bz-NCS dissolved in DMSO in 0.1 M sodium carbonate buffer (pH 9) at 37°C for 1 h. The resulting trastuzumab-DFO conjugate was purified via a gel filtration spin column (molecular weight cut off, 40 kDa) during which the buffer was exchanged to 1 M 4-(2-hydroxyethyl)-1-piperazineethanesulfonic acid (HEPES) buffer. The product was then sterile-filtered into 0.5-mL doses in a laminar flow hood and stored in a freezer at -80˚C. Production of $^{89}$Zr-oxalate was acquired for 3D Imaging (Little Rock, AR) diluted with an equal volume of 1 M HEPES buffer and neutralized to a pH range previously reported for optimal radiolabeling (pH 6.8‑7.2) with 2 M sodium hydroxide and 2 M hydrochloric acid. The reaction solution consisting of trastuzumab-DFO (10 mg/mL), $^{89}$Zr-oxalate (370 MBq/mL), and 1 M HEPES buffer was pumped into a single channel reactor (total volume, ~2.81 μL) from separate inlets in varying ratios of ligand to metal at a total flow rate of 20 μL/min. Ratios of mg trastuzumab-DFO to MBq of $^{89}$Zr greater than 1:148 were incubated for 1 h at 37° C within the reactor (by halting the flow), whereas lower ratios of mAb:$^{89}$Zr were pumped through at a continuous rate. After the reaction, the reactor was flushed with 1 M HEPES buffer to wash out the remaining $^{89}$Zr-trastuzumab; the product was collected in a microcentrifuge tube and the radiochemical yield was confirmed by instant thin-layer chromatography.

*Radiosynthesis of [$^{18}$F] Fluoro-2-deoxy-D-glucose:*

2-[$^{18}$F]fluoro-2-deoxy-D-glucose (FDG) production was manufactured at UCSF on a GE FASTlabTM synthesizer using an FDG citrate disposable cassette. Approximately 7 Ci of [$^{18}$F] was delivered to the synthesizer after the bombardment of [$^{18}$O]-water with hydrogen. $^{18}$F- was trapped on a quaternary methyl ammonium (QMA) anion exchange cartridge and eluted with a 0.5 mL solution of Kryptofix (K2.2.2), potassium carbonate, acetonitrile (MeCN), and water solution. Acetonitrile additions form an azeotropic mixture for evaporation of residual water. Mannose triflate precursor dissolved in MeCN was added to the reaction vessel for a 3-minute labelling reaction at 125˚C. The resulting FTAG (2-[$^{18}$F]-fluoro-1,3,4,6-tetra-O-acetyl-D-glucose was then diluted with water prior to transfer to the reversed-phase tC18 cartridge. FTAG was retained on the tC18 while unreacted $^{18}$F and impurities flow through the cartridge to waste. Alkaline hydrolysis of the trapped FTAG on the tC18 was performed with 2N NaOH. FDG was eluted with water and neutralized in 3.2 mL of citrate buffer before final purification through a tC18 plus cartridge and Alumina N cartridge. Following this 25-minute synthesis, the final product was delivered through a 0.22μm vented filter to the final product vial. The synthesis resulted in an approximately 80% decay-corrected yield.

**Supplemental Figures:**

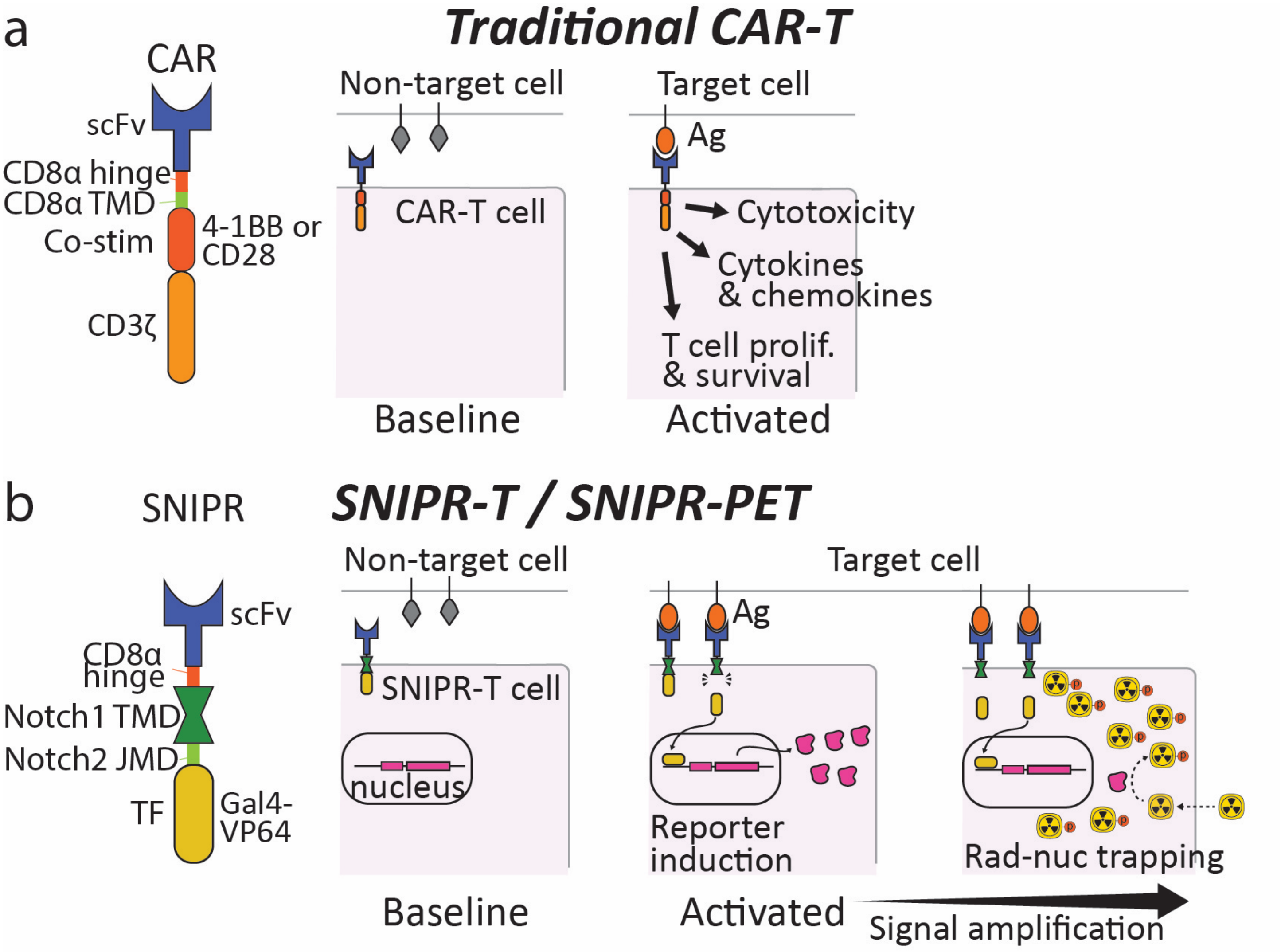

**Supplemental Figure 1: SNIPR-T cells and SNIPR-PET, in comparison to traditional chimeric antigen rector (CAR)-T cells.** A. CAR comprises an extracellular antigen recognition domain (scFv), CD8α hinge domain and CD8α transmembrane domain (TMD), co-stimulatory domains (Co-stim) such as 4-1BB or CC28 and a CD3ζ domain. When scFv of CAR binds to its target antigen, intracellular domains trigger a native T cell response. The native T cell response includes cytotoxicity, cytokines, chemokine and growth factor secretion, and T cell proliferation and survival. B. SNIPR consists of an extracellular antigen recognition domain (scFv), CD8α hinge domain, Notch1 TMD, Notch2 Juxtamembrane domain (JMD) and transcription factor (TF) domain Gal4. When scFv of SNIPR binds to its target antigen, TMD is cleaved and TF falls off from the receptor, entering the nucleus to activate pre-programmed gene expression.

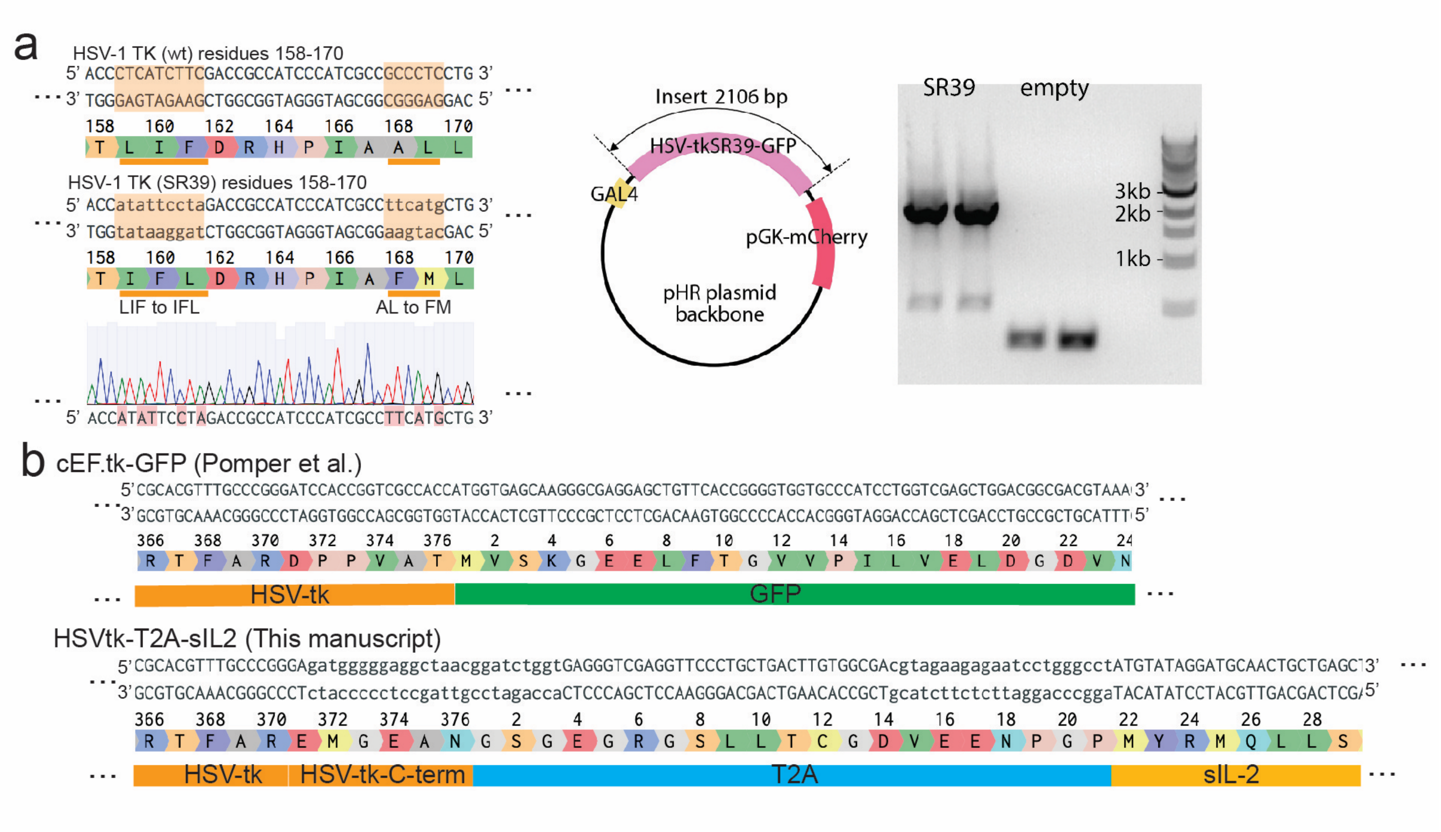

**Supplemental Figure 2: Cloning of HSV-tkSR39.** A. We cloned HSV-tk (SR39) from cEF.tk-GFP[4] from Addgene plasmid depositary as a template (left). Point mutations were generated by using primers GGCGATGGGATGGCGGTCtAgGAatAtGGTGAGGGCCGGGGGC and GACCGCCATCCCATCGCCttCaTgCTGTGCTACCCGGCC, followed by In-Fusion cloning (Takara Bio) into the pHR plasmid with pGK-mCherry. pGK-mCherry was used for selecting T cell containing this plasmid (middle). PCR and DNA electrophoresis was used to confirm the plasmid. B. 6 amino acids C-terminal of HSV-tk was truncated modified in the original clone (cEF.tk-GFP), therefore added to match the HSV-tk sequence from UniProt (Wagner et al., PMID: 6262799) using primer gTCGCCACAAGTCAGCAGGGAACCTCGACCCTCaccagatccgttagcctcccccatcTCCCGGGCAAACGTGC and ctctcgacattcgttggatc. This PCR product was cloned with T2A and super IL2 (sIL-2) for co-induction of HSV-tk reporter and sIL-2.

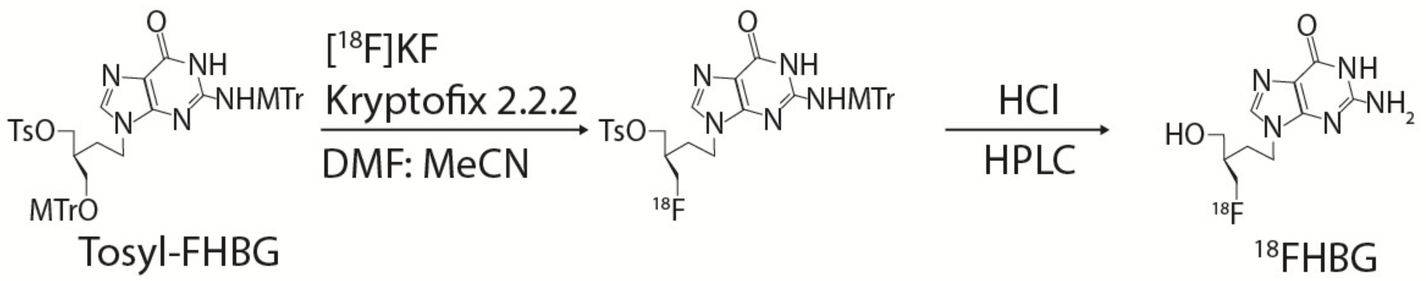

**Supplemental Figure 3: Radiosynthesis of [$^{18}$F]FHBG.** The [$^{18}$F]fluoride ion was trapped on a QMA cartridge (Waters, MA) and eluted with $K_2CO_3$/ K2.2.2 in acetonitrile/water. The resultant solution was azeotroped at 110˚C twice with additions of acetonitrile. To this dried vial, the precursor N2, monomethoxytrityl-9-[4-(tosyl)-3-monomethoxytrityl-methylbutyl] guanine (or Tosyl-FHBG) (ABX, Radeberg, Germany) dissolved in acetonitrile (0.8 mL) was added and heated to 140 ˚C for 10 minutes. Then 1 mL of 1M HCl was added and the reactor was heated to 105 ˚C for 5 minutes. The reaction was cooled, neutralized with 0.5 mL of 2 M NaOH, diluted with solvent system (2 mL), and injected onto the semi-preparative HPLC to collect, filtered and analyze dvia HPLC.

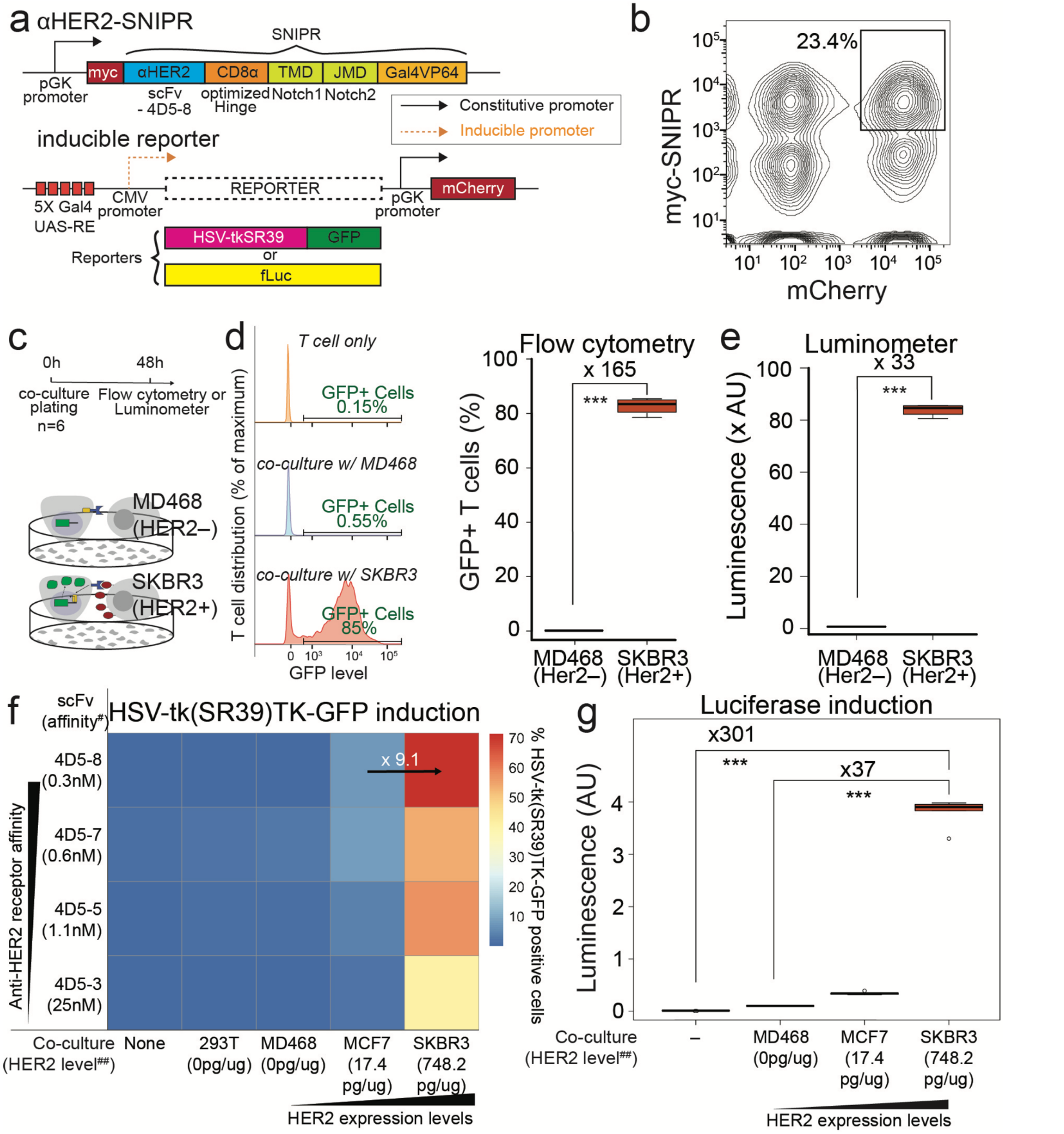

**Supplemental Figure 4: Anti-HER2-SNIPR activation and reporter expression.** A. To demonstrate the feasibility of SNIPR-PET, we engineered T cells capable of SNIPR-induced thymidine kinase expression and SNIPR-induced luciferase enzymatic activity. We generated cells bearing an anti-HER2 SNIPR and inducible HSV-tkSR39-GFP or inducible luciferase reporters. The reporter plasmids contain constitutively expressed mCherry fluorescent protein, which is also used for cell sorting. SNIPR T cells are generated by double transduction using lentivirus. B. T cells expressing both SNIPR and mCherry are sorted using fluorescence activated cell sorting (FACS). The double positive T cells were about 15-50%. C. SNIPR T cells are co-cultured with either MD468 (HER2– cells, negative control) or SKBR3 (HER2+ cells) for 48 hours. D. TK-GFP reporter expression is detected using FAC analysis. An example of FAC analysis for TK-GFP expression is shown, in anti-HER2 SNIPR T cells only (top, left), anti-HER2 SNIPR T cells with HER2- cells (middle, left) and anti-HER2 SNIPR T cells with HER2+ cells (bottom, left). There was an over 150-fold increase in number of HSV-tkSR39-GFP+ cells after incubating with SKBR3 (HER2+) compared to after incubating with MD468 (HER2-)(right). E. Activation of antiHER2-SNIPR induced an over 33-fold increase in luciferase enzymatic activity after incubating with SKBR3 (HER2+) compared to after incubating with MD468 (HER2-). F. We tested the effect of receptor binding affinity and target antigen abundance to HSV-tkSR39-GFP induction by using anti-HER2 SNIPR cells with varying HER2 binding affinities, and by using cancer cells with varying levels of HER2 expression. As expected, increasing the HER2 binding affinity of SNIPR receptor and increasing HER2 abundance resulted in higher induction of HSV-tkSR39-GFP. G. We tested the effect of increasing target antigen abundance to the SNIPR-induced luciferase enzymatic activity. Increasing target antigen abundance demonstrated a higher level of luciferase enzymatic activity. *$p<0.05$, **$p<0.01$, ***$p<0.001$. #scFv affinity values from Zhao et al.[10], ##HER2 abundance values from Collins et al.[11].

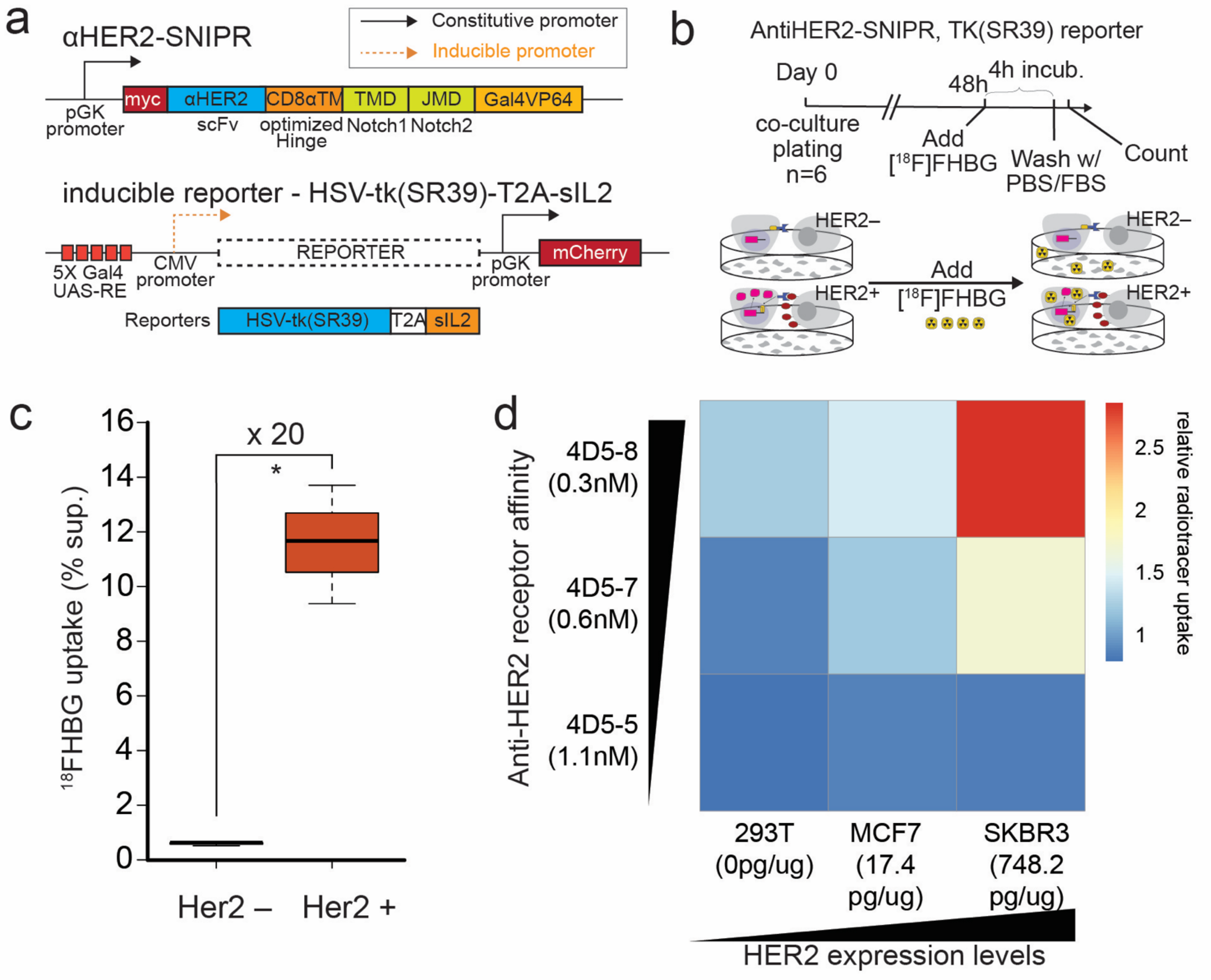

**Supplemental Figure 5. *In vitro* [$^{18}$F]FHBG uptake in activated anti-HER2 SNIPR T cells.** A. Constructs that were used for the *in vitro* SNIPR-induced [$^{18}$F]FHBG uptake experiment. We generated T cells bearing an anti-HER2 SNIPR receptor and inducible HSV-tk(SR39)-T2A-sIL2, followed by FACS with myc and mCherry. B. Anti-HER2 SNIPR T cells were co-cultured with HER2+ (SKBR3) or HER2- (MD468) cells for 48 hours, followed by 3 hour incubation with [$^{18}$F]FHBG. C. SNIPR T cells co-cultured with HER2+ (SKBR3) cells accumulated significantly higher amount of [$^{18}$F]FHBG radiotracer compared to T cells co-cultured with HER2– (MD468) cells. D. Similar to **Fig. 1F**, we tested the effect of receptor binding affinity and target antigen abundance on [$^{18}$F]FHBG accumulation by using anti-HER2 SNIPR with varying HER2 binding affinities, and by using cancer cells with varying levels of HER2 expression. As expected, increasing HER2 binding affinity of SNIPR receptor and increasing HER2 abundance resulted in greater enrichment of [$^{18}$F]FHBG. *$p<0.05$, **$p<0.01$, ***$p<0.001$.

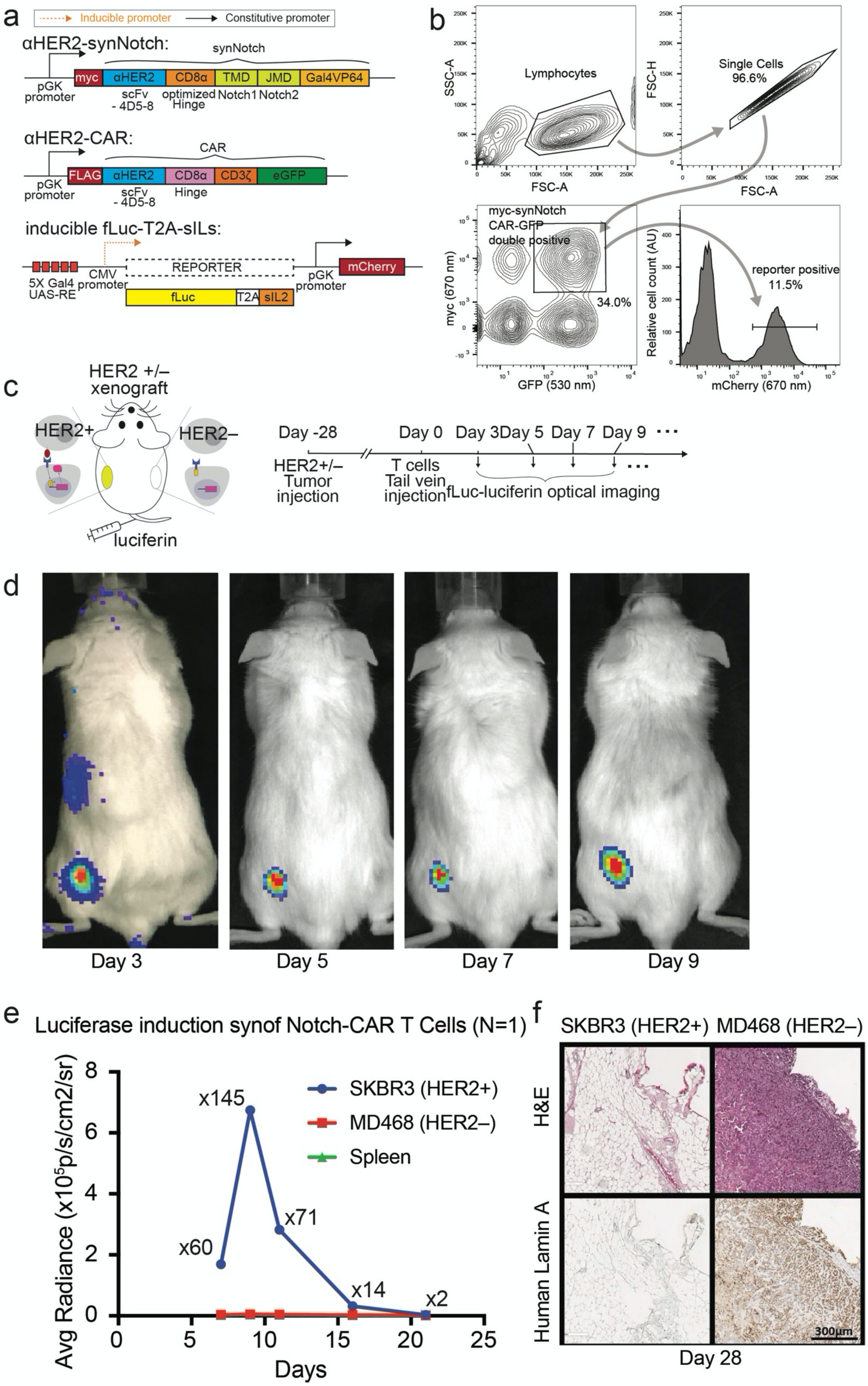
a
Inducible promoter
Constitutive promoter
αHER2-synNotch:
synNotch
pGK promoter
myc
αHER2
CD8α
TMD
JMD
Gal4VP64
scFv - 4D5-8
optimized Hinge
Notch1
Notch2
αHER2-CAR:
CAR
FLAG
CD3ζ
eGFP
Hinge
inducible fLuc-T2A-sILs:
5X Gal4 UAS-RE
CMV promoter
REPORTER
fLuc
T2A
sIL2
mCherry
b
Lymphocytes
SSC-A
FSC-A
Single Cells 96.6%
FSC-H
myc-synNotch CAR-GFP double positive
34.0%
myc (670 nm)
GFP (530 nm)
reporter positive 11.5%
Relative cell count (AU)
mCherry (670 nm)
c
HER2 +/– xenograft
HER2+
HER2–
luciferin
Day -28
Day 0
Day 3
Day 5
Day 7
Day 9
HER2+/– Tumor injection
T cells Tail vein injection
fLuc-luciferin optical imaging
d
Day 3
Day 5
Day 7
Day 9
e
Luciferase induction synof Notch-CAR T Cells (N=1)
Avg Radiance (x10⁵p/s/cm2/sr)
Days
x60
x145
x71
x14
x2
SKBR3 (HER2+)
MD468 (HER2–)
Spleen
f
SKBR3 (HER2+) MD468 (HER2–)
H&E
Human Lamin A
300μm
Day 28

**Supplemental Figure 6: Effect of CAR to *in vivo* luciferase imaging.** A. Anti-HER2 T cells bearing anti-HER2 synNotch, anti-HER2-CAR and the inducible luciferase-T2A-sIL2 reporter were generated. B. Lentivirally transduced T cells were sorted based on myc on anti-HER2 synNotch, GFP on anti-HER2 CAR and mCherry on the reporter construct, yielding 8-12% triple positive T cells. C. Dual xenograft was generated by injecting human HER2+ and HER2– cancer cell lines 4 weeks before synNotch T cell injection. Luciferase imaging was acquired up to 21 days after injecting luciferin. D. Representative bioluminescence imaging demonstrates strong signal only within the HER2+ tumors. E. Signal within HER2+ tumor increased up to day 9 and decreased afterwards, likely secondary to weak target killing activity of CD4+ T cells with CAR activation. F. At day 28 after T cell injection, mice were sacrificed for histologic analysis. There was no human lamin A + cells at HER2+ cells, suggesting clearance of HER2+ tumor by synNotch-CAR T cells.

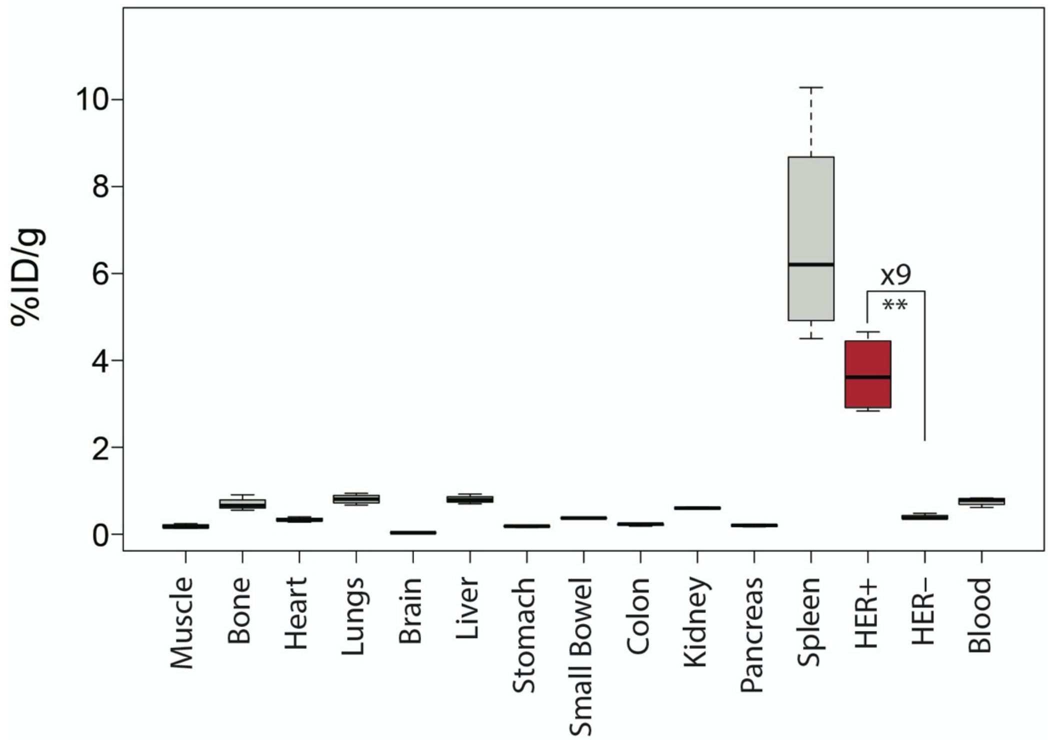

**Supplemental Figure 7: Biodistribution analysis of [$^{89}$Zr]Herceptin at day 3.** Biodistribution analysis of [$^{89}$Zr]Herceptin at day 3 demonstrated significant greater [$^{89}$Zr]Herceptin enrichment within HER2+ xenograft compared to HER2– xenograft (n=4, p=0.005). As seen on microPET-CT, the spleen demonstrated a high level of [$^{89}$Zr]Herceptin uptake.
